# Peptidergic modulation of fear responses by the Edinger-Westphal nucleus

**DOI:** 10.1101/2021.08.05.455317

**Authors:** Michael F. Priest, Sara N. Freda, Deanna Badong, Vasin Dumrongprechachan, Yevgenia Kozorovitskiy

## Abstract

Many neuronal populations that release fast-acting excitatory and inhibitory neurotransmitters in the brain also contain slower acting neuropeptides. These facultative peptidergic cell types are common, but it remains uncertain whether obligate peptidergic neurons exist. Our fluorescence in situ hybridization, genetically-targeted electron microscopy, and electrophysiological characterization data strongly suggest that neurons of the non-cholinergic, centrally-projecting Edinger-Westphal nucleus in mice are fundamentally obligately peptidergic. We further show, using fiber photometry, monosynaptic retrograde tracing, anterograde projection mapping, and a battery of behavioral assays, that this peptidergic population both promotes fear responses and analgesia and activates in response to loss of motor control and pain. Together, these findings elucidate an integrative, ethologically relevant function for the Edinger-Westphal nucleus and functionally align the nucleus with the periaqueductal gray, where it resides. This work advances our understanding of the peptidergic modulation of fear and provides a framework for future investigations of putative obligate peptidergic systems.

## Introduction

Numerous glutamatergic and GABAergic neurons in the mammalian brain have been shown to be facultatively peptidergic (Dicken et al., 2012; Hökfelt et al., 2018; Merighi et al., 2011; Nusbaum et al., 2017; Van Den Pol, 2012; Schöne and Burdakov, 2012). That is, they can package and release one or more neuropeptides, but the neurons also release fast-acting neurotransmitters independently of peptide release. Similar phenomena are observed in other classes of neurons that release fast neurotransmitters with small molecule neuromodulators including biogenic amines or acetylcholine (Granger et al., 2020; Hnasko et al., 2010; Stuber et al., 2010; Tritsch et al., 2012; Vaaga et al., 2014; Wang et al., 2019; Zhang et al., 2015), or release biogenic amines as well as neuropeptides (Hökfelt et al., 2018; Merighi et al., 2011; Nusbaum et al., 2017; Van Den Pol, 2012). Different chemical signaling molecules released from the same neuronal population can produce distinct behavioral responses (Pomrenze et al., 2019). Furthermore, neuropeptides and neurotransmitters can use different machinery for packaging and release, e.g., large dense core vesicles versus small clear vesicles (Hökfelt et al., 2018; Merighi et al., 2011; Van Den Pol, 2012; Schöne and Burdakov, 2012). However, it is not known whether obligate peptidergic neurons, which release no fast-acting canonical neurotransmitters, exist. Indeed, recent reviews have been skeptical of the existence of obligate peptidergic neurons (Hökfelt et al., 2018; Van Den Pol, 2012), or have suggested that this class may be limited to the neurosecretory cells of the paraventricular nucleus of the hypothalamus (PVN) (Merighi et al., 2011).

Recent single cell RNA-sequencing (scRNA-seq) of the mouse brain has pointed to the possible existence of obligate peptidergic neuronal populations both within and outside the PVN (Huang et al., 2019; Romanov et al., 2017; Zeisel et al., 2018). Due to technical limitations associated with sequencing depth in scRNA-seq, the absence of a given transcript from a cell type does not preclude its existence. Nevertheless, scRNA-seq data highlight several neuronal populations that represent obligate peptidergic candidates (Figure 1A). One potential candidate is the cocaine-and-amphetamine regulated transcript (CART)-positive neuronal population of the Edinger-Westphal (EW) nucleus.

**Figure 1:**
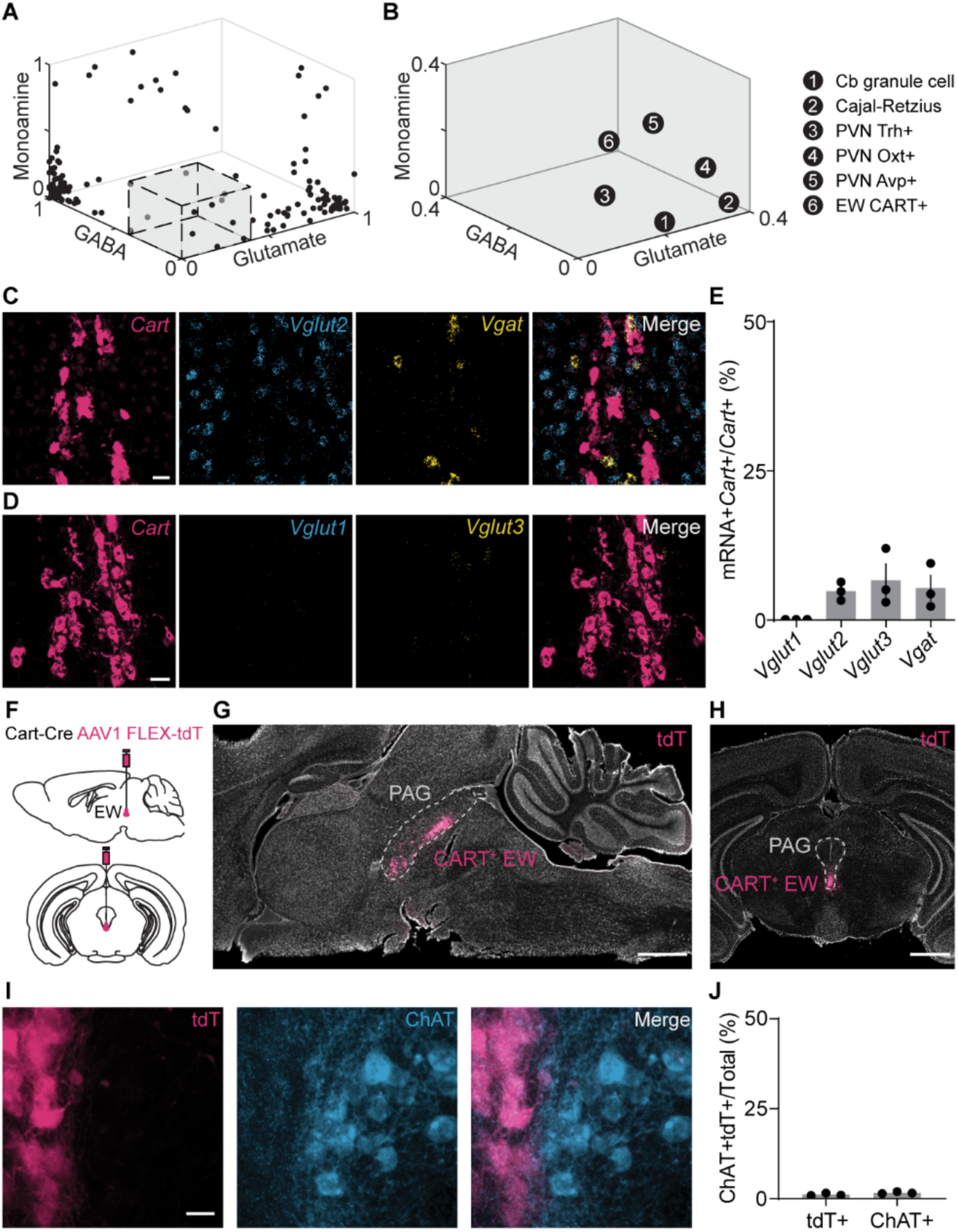
The CART^+^ EW lacks molecules required for fast neurotransmission. (A) Neuronal types in the mouse brain, by the proportion of cells in each group that contain markers for GABA, glutamate, and monoamines inclusive of acetylcholine. Many neuronal types have high proportions of cells that contain markers of inhibitory or excitatory neurons. The gray box, enlarged in (B), delineates neuronal types with very few cells identified as glutamatergic, GABAergic, or monoaminergic. Data are reanalyzed from publicly available scRNA datasets (Zeisel et al., 2018). (B) As in (A), but for neuronal types with <0.4 of any neurotransmitter or neuromodulator marker. The 6 neuronal types: cerebellar granule cells, Cajal-Retzius cells, Trh^+^, Oxt^+^, and Avp^+^ neurons of the hypothalamic periventricular nucleus, and CART^+^ neurons of the Edinger-Westphal nucleus. (C) Example FISH of mRNA encoding CART (red, *Cartpt* or *Cart*), Vglut2 (blue, *Slc17a6* or *Vglut2*), and Vgat (yellow, *Slc32a1* or *Vgat*) in the EW. Scale bar, 20 µm. (D) As in (C), but against *Vglut1* (blue, *Slc17a7* or *Vglut1*), and *Vglut3* (yellow, *Slc17a8* or *Vglut3*). (E) Colocalization of vesicular transporters with *Cart* in the EW (*Vglut1*, 0.00% ± 0.00%; *Vglut2*, 4.83% ± 0.90%; *Vglut3*, 6.70% ± 2.72%; *Vgat*, 5.40% ± 2.14%; n= 3 mice for all; 297 CART^+^ cells for *Vglut2* and *Vgat*; 404 CART^+^ cells for *Vglut1* and *Vglut3*). (F) Schematic of AAV1 FLEX-tdTomato (tdT) injection into the mouse CART-Cre EW. (G) Selective tdT (red) expression in the CART^+^ EW, sagittal slice. The PAG is marked by the dashed gray line. Hoechst nuclear stain, white. Scale, 1 mm. (H) As in (G), but for a midbrain coronal slice. (I) tdT+ neurons of the EW (red) compared to immunofluorescence against ChAT (blue). Scale, 20 µm. (J) CART^+^ EW neurons do not colocalize with ChAT^+^ neurons (ChAT^+^CART^+^/CART^+^, 1.1% ± 0.2%, n = 3 mice, 784 cells; ChAT^+^CART^+^/ChAT^+^, 1.7%±0.2%, n=3 mice, 879 cells). Error bars represent SEM. See also Figure S1.

The CART^+^ EW resides in the rostral ventromedial periaqueductal gray, aligning with the anatomical regions labeled as Edinger-Westphal and pre-Edinger-Westphal nucleus in rodent brain atlases (Lein et al., 2007; Paxinos and Franklin, 2019). The CART^+^ EW contains multiple neuropeptides including CART, urocortin (Ucn), and cholecystokinin (CCK), and it projects to the spinal cord (Kozicz et al., 2011). These properties distinguish the CART^+^ EW, also known as the centrally-projecting EW, from both the cholinergic EW (Vasconcelos et al., 2003) that projects to the ciliary ganglion and controls lens accommodation (Toyoshima et al., 1980) and from the glutamatergic CCK^+^ peri-EW neuronal population that promotes non-REM sleep and lacks the descending projections (Zhang et al., 2019) that are characteristic of CART^+^ EW neurons.

The CART^+^ EW was recently classified as peptidergic, although not obligately peptidergic, in a scRNA-seq study of the mouse dorsal raphe (Huang et al., 2019). In contrast, another scRNA-seq study of cells from across the entire mouse brain classified the CART^+^ EW as glutamatergic (Zeisel et al., 2018). In addition, prior optogenetic experiments showed light-evoked glutamate-mediated excitatory postsynaptic currents (EPSCs) in the medial prefrontal cortex following Cre-dependent viral vector-driven channelrhodopsin (ChR2) expression in CCK-Cre^+^ neurons of the EW, which colocalize with CART (Li et al., 2018). To resolve this discrepancy, we combined fluorescence in situ hybridization, genetically-targeted electron microscopy, whole central nervous system anterograde mapping, and electrophysiological characterization to assess whether the CART^+^ EW is an obligate peptidergic nucleus.

Other candidate obligate peptidergic neurons in the PVN have been shown to modulate fear or anxiety (Hasan et al., 2019; Knobloch et al., 2012; Rigney et al., 2021), and the CART^+^ EW responds to stressors (Gaszner et al., 2004) and contains neuropeptides that alter fear and anxiety (Bowers et al., 2012; Stanek, 2006; Sztainberg and Chen, 2012). Therefore, we used chemogenetic modulation and genetically-targeted ablation to test whether the CART^+^ EW mediates fear responses and used monosynaptic retrograde labeling and *in vivo* calcium fiber photometry to interrogate the circuits and stimuli that regulate the CART^+^ EW. Together, our data define the CART^+^ EW as an obligate peptidergic nucleus that promotes fear responses.

## Results

### The CART^+^ EW lacks molecules required for fast neurotransmission

We defined candidate obligate peptidergic nuclei using an unbiased scRNA-seq study of the mouse nervous system that identified 181 neuronal clusters, or cell types, in the central nervous system (Zeisel et al., 2018). We excluded neuroblast-like cell types and spinal cord cells, focusing on the proportion of cells in each of 148 neuronal brain cell types that contained canonical markers of glutamate, GABA/glycine, monoamines or acetylcholine (Figure 1A). For the remainder of this paper, we will group acetylcholine, which is a monoammonium, with the canonical monoamines: serotonin, dopamine, epinephrine, and norepinephrine. Additionally, although glutamate, GABA, and glycine are chemically monoamines, we will categorize glutamate and GABA/glycine separately. Glutamatergic markers were the vesicular glutamate transporters *Slc17a6, Slc17a7,* and *Slc17a8*. GABAergic markers were the vesicular GABA transporter, *Slc32a1*, the glycine transporters *Slc6a5* and *Slc6a9*, and the GABA-synthesizing glutamate decarboxylase enzymes *Gad1* and *Gad2*. Monoaminergic markers were the transporters *Slc5a7*, *Slc6a3*, *Slc6a4*, *Slc18a1*, *Slc18a2*, and *Slc18a3*, biosynthetic enzymes *Chat, Dbh, Ddc, Hdc, Pnmt*, *Th*, *Tph1*, and *Tph2*, and the serotonergic cell marker *Fev*. For each cell, the presence of any amount of transcript was used to identify a cell as glutamatergic, GABAergic, and/or monoaminergic, and any cell could be positive for all three categories. As expected, many neuronal types previously defined by clustering (Zeisel et al., 2018) had large proportions of cells that contained markers for glutamate, GABA, or a monoamine (Figure 1A).

To find candidates for obligate peptidergic cell types, we focused on cell types with a minority (<40%) of their cells positive for any of the 23 markers (Figure 1B). Only six neuronal types fit these criteria: the Cajal-Retzius cell of the hippocampus, the granule cells of the cerebellum, the vasopressin-positive (Avp^+^), oxytocin-positive (Oxt^+^), and thyrotropin-releasing hormone-positive (Trh^+^) neurons of the PVN, and the CART^+^ neurons of the Edinger-Westphal nucleus. It is unclear why granule cells of the cerebellum appear here, as they are known to be glutamatergic, but their extremely small size (Lackey et al., 2018) likely contributes to technical difficulties in sequencing depth (Zeisel et al., 2018); their inclusion underscores the importance of validating scRNA-seq results of interest with lower throughput approaches. Cajal-Retzius cells are crucial for proper cortical development in early life (Ogawa et al., 1995) and may have a predominantly developmental role, rather than one dependent on neurotransmission in established circuits (Causeret et al., 2021). The neurohypophysiotrophic populations of the PVN have been previously suggested as candidate obligate neuropeptidergic populations (Merighi et al., 2011).

The remaining candidate is the CART^+^ population of the EW, which was found to have glutamatergic markers in 17.0% of its 47 cells, GABAergic markers in 12.8% of cells, and monoaminergic or cholinergic markers in 23.4% of cells. Only 4.3% of the CART^+^ EW contained both a biosynthetic enzyme and a related transporter, *e.g*., tyrosine hydroxylase with a vesicular monoamine transporter. Another scRNA-seq study also found that the CART^+^ EW lacked monoaminergic or cholinergic markers (Huang et al., 2019), and the CART^+^ EW is not considered a monoaminergic or cholinergic population (Kozicz et al., 2011). However, despite apparently lacking mRNA encoding the transporters required for vesicular packaging, one recent report has claimed that the CART^+^ EW is glutamatergic (Li et al., 2018).

One possibility is that scRNA-seq failed to pick up transcripts for vesicular glutamate or GABA transporters. We tested the presence of these transporters in the CART^+^ EW using fluorescence *in situ* hybridization (FISH) against mRNAs for CART (*Cartpt* or *Cart*) and vesicular transporters for glutamate (*Slc17a6*/*Vglut2*, *Slc17a7*/*Vglut1*, *Slc17a8*/*Vglut3*) and GABA (*Slc32a1*/*Vgat*) (Figures 1C and 1D). Male and female mice were included in all experiments. The vast majority of CART^+^ EW neurons lack canonical transporters for glutamate (*Vglut1*, 0.00% ± 0.00%; *Vglut2*, 4.83% ± 0.90%; *Vglut3*, 6.70% ± 2.72%, n = 3 mice, 404 *Cart*^+^ cells for *Vglut1* and *Vglut3*, 297 *Cart*^+^ cells for *Vglut2*) or GABA (*Vgat*, 5.40% ± 2.14%, n = 3 mice, 297 *Cart*^+^ cells) (Figures 1C-1E and S1A-D). This FISH quantification shows that approximately 83% of CART^+^ EW neurons in mice lack any canonical vesicular transporter for glutamate, GABA, or glycine release, suggesting that much of the CART^+^ EW may function as an obligate peptidergic neuronal population.

The absence of *Vglut1*^+^ cells in subcortical areas was contrasted by hippocampal neurons in the same tissue, which expressed *Vglut1* robustly (Figure S1E). Immediately adjacent to and within the Edinger-Westphal nucleus, we observed numerous CART-negative *Vglut2*^+^ (n = 771 cells, 3 mice) and *Vgat*^+^ (n = 256 cells, 3 mice) neurons, as well as fewer *Vglut3*^+^ neurons (n = 11 cells, 3 mice), underscoring the necessity of distinguishing the genetically-defined CART^+^ EW population from the anatomically-defined EW.

Therefore, to genetically target the CART^+^ EW, we used the Cart-IRES2-Cre mouse line (Cart-Cre) for experiments throughout the study (Daigle et al., 2018). Following injection of adeno-associated viral vector (AAV) packaged with a Cre-dependent tdTomato (tdT) gene into the Edinger-Westphal nucleus (Figure 1F), we observed robust tdT expression that was limited to cells in this region (Figures 1G and 1H). Quantification of tdT^+^ cells revealed that the CART^+^ EW is comprised of around 1,200 Cart-Cre neurons (1189 cells ± 90.6, n = 3 mice). As expected, genetically-defined CART^+^ EW shows virtually no overlap (1.15% ± 0.21%, n = 3 mice, 1185 cells) with preganglionic, cholinergic neurons of the EW or other nearby oculomotor nuclei labeled with choline acetyltransferase (ChAT) (Figures 1I and 1J) (Vasconcelos et al., 2003; Weitemier et al., 2005). This confirms that the genetically-targetable CART^+^ EW lacks the required biosynthetic enzyme for producing the fast-acting neurotransmitter acetylcholine. We also confirmed that the AAV-targeted CART^+^ EW colocalizes heavily with immunolabeled CART (85.12% ± 1.52%, n = 3 mice, 784 cells) (Lovett-Barron et al., 2017) and urocortin (80.01% ± 1.44%, n = 3 mice, 631 cells) (Figure S2A to S2D). The CART and urocortin populations of the EW appear to be largely identical, consistent with prior work (Kozicz, 2003; Xu et al., 2009).

### The CART^+^ EW contains numerous neuropeptides and large vesicles

In addition to CART and urocortin, the EW has been suggested to release other neuropeptides. To refine our understanding of peptidergic motifs in the EW, we examined the colocalization of EW *Cart* with substance P (*Tac1*) (Phipps et al., 1983; Skirboll et al., 1982), cholecystokinin (*Cck*) (Huang et al., 2019; Li et al., 2018; Maciewicz et al., 1984; Zeisel et al., 2018), pituitary adenylate cyclase-activating peptide (*Adcyap1*, or *Pacap*), and neuromedin B (*Nmb*) (Burnell et al., 2008) (Figures 2A, S2E, and S2F). We found no evidence for co-expression of *Cart* with *Tac1* (2.86% ± 0.92%, n = 2 mice) in contrast to *Cck* (74.04% ± 2.34%, n = 3), *Nmb* (67.17% ± 6.28%, n = 3), and *Pacap* (54.12% ± 6.16%, n = 3) (Figures 2A, 2B, S2E, and S2F). Thus, the CART^+^ EW contains numerous neuropeptides, including CART, urocortin, CCK, neuromedin B, and PACAP, but lacks molecules required for fast neurotransmission, including glutamate and GABA vesicular transporters as well as ChAT.

**Figure 2:**
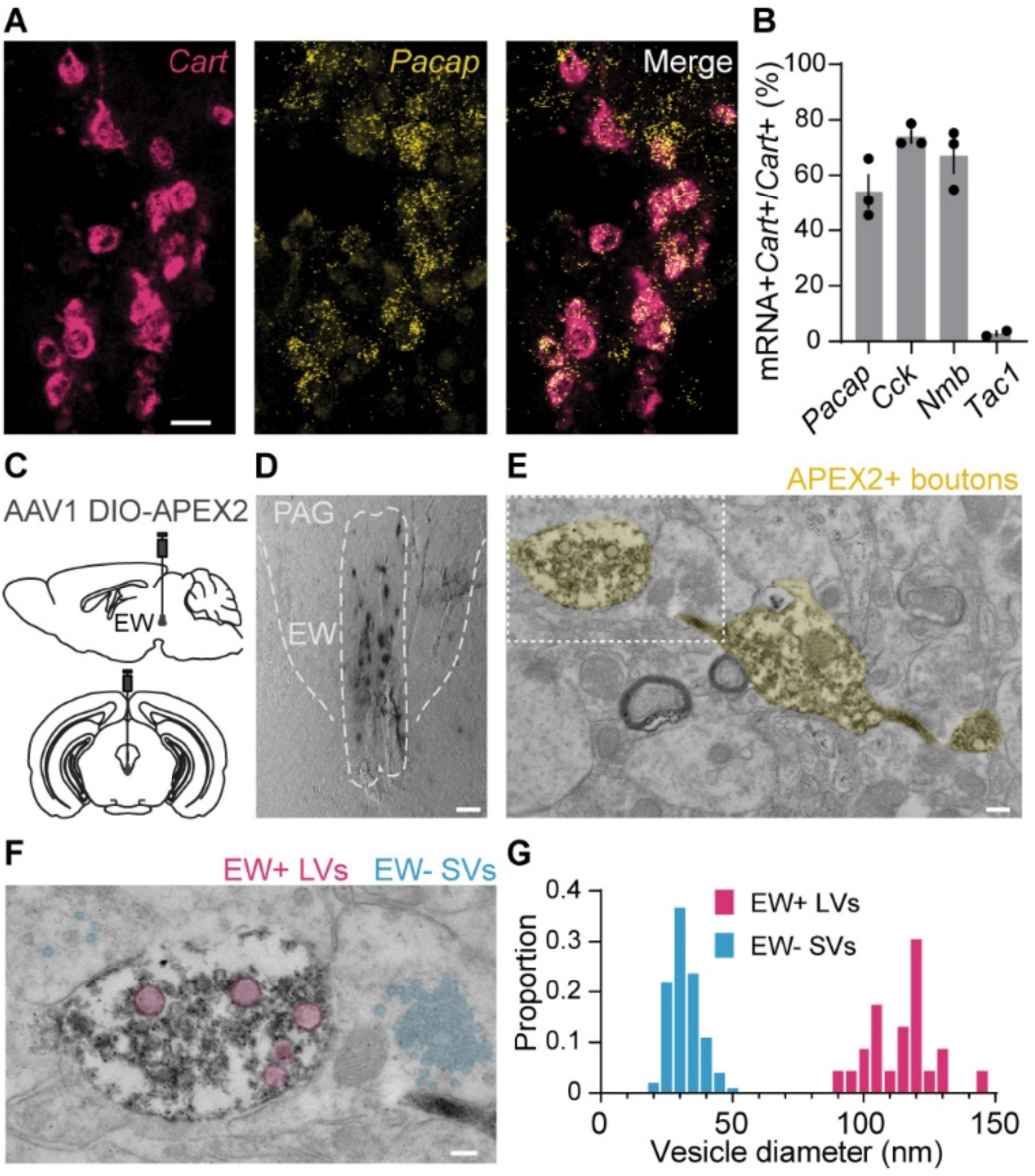
The CART^+^ EW contains numerous neuropeptides and large vesicles. (A) FISH against *Cart* (red) and *Adcyap1*/*Pacap* (yellow). Scale, 20 µm. (B) Quantification of *Cart* colocalization with *Pacap*, *Cck*, *Nmb*, and *Tac1*. *Pacap*, 54.12% ± 6.16%; *Cck*, 74.04% ± 2.34%; *Nmb*, 67.17% ± 6.28%; *Tac1*, 2.86% ± 0.92%; n= 2 mice for *Tac1*, 3 mice for all others. (C) Schematic of AAV1 DIO-APEX2 injection into the mouse CART-Cre EW. (D) Brightfield image of genetically targeted APEX DAB staining of CART^+^ EW cells. Scale, 50 μm. (E) Transmission electron micrograph of APEX2^+^ synaptic boutons in the midbrain. Scale, 200 nm. (F) Higher magnification image of the white box in (E). Multiple small clear vesicles (blue) are observed in synaptic processes around the APEX2^+^ bouton, which contains multiple large vesicles (red). Scale, 100 nm. (G) Diameters of dense core vesicles (red) observed in APEX^+^ boutons and small clear vesicles (blue) found in regions adjacent to the APEX2^+^ boutons; n = 23 large vesicles from 16 APEX2^+^ boutons, diameter = 114.1 nm ± 2.6 nm; n = 101 small vesicles near 9 APEX2^+^ boutons, diameter = 31.8 nm ± 0.6 nm. Histogram bin size is 5 nm. Error bars represent SEM. See also Figure S2.

Next, we targeted the CART^+^ EW with Cre-dependent LCK-APEX2, an engineered peroxidase (APEX) capable of performing proximity labeling and depositing electron dense diaminobenzidine. APEX is fused to a membrane anchor sequence (LCK) for membranal localization with a separately transcribed GFP for identifying expression (Dumrongprechachan et al., 2021). Following APEX expression in the CART^+^ EW, fixed, stained slices were imaged under a transmission electron microscope. In addition to labeling structures in the soma, APEX^+^ boutons were abundant in the imaged peri-EW midbrain regions. These boutons contained numerous large vesicles (n = 23 vesicles from 16 boutons, diameter = 114.1 nm ± 2.6 nm) matching the size of large dense core vesicles (Hökfelt et al., 2018). No small clear vesicles were observed in these boutons, although nearby structures frequently contained vesicles the size of small clear vesicles (n = 101 vesicles near 9 APEX^+^ boutons, diameter = 31.8 nm ± 0.6 nm). We cannot preclude the possibility that our sample preparation may have obscured the presence of small clear vesicles in the APEX^+^ boutons. However, the marked presence in CART^+^ EW boutons of the large vesicles necessary for peptidergic release and the apparent absence of the small clear vesicles used for fast neurotransmission (Hökfelt et al., 2018; Merighi et al., 2011; Van Den Pol, 2012; Schöne and Burdakov, 2012) supports the hypothesis that the CART^+^ EW is obligately peptidergic.

### The CART^+^ EW projects to the spinal cord and multiple subcortical regions

To functionally assess whether a population is obligately or facultatively peptidergic, it is important to know its efferent targets. However, the targets of EW projections within the brain (Li et al., 2018; Dos Santos Júnior et al., 2015) and spinal cord (Bittencourt et al., 1999; Loewy and Saper, 1978; Dos Santos Júnior et al., 2015) remain disputed. To resolve these conflicting reports and enable functional characterization of CART^+^ EW signaling, we mapped the CART^+^ EW anterograde projections using genetically-targeted AAV tdT injections into the EW of CART-Cre mice, with subsequent immunofluorescence against tdT (Figure 3A). We found robust projections to laminae III, IV, and X of the spinal cord (Figure 3B, n = 3 mice) using isolectin B4 labeling (Figure S3A). We registered brain sections (Figure 3C) to the Allen Brain Atlas for automated quantification of CART^+^ EW projections in each brain region (Figures 3D, 3E, S3B, and S3C, n = 3 mice). Projections were predominantly found in numerous subcortical brain regions. This finding partially aligns with non-genetically defined anterograde tracing of the rat anatomical EW (Dos Santos Júnior et al., 2015), but contradicts genetically defined transsynaptic viral expression from a CCK^+^ EW population, which found target neurons primarily in the medial prefrontal and cingulate cortices (Li et al., 2018).

**Figure 3:**
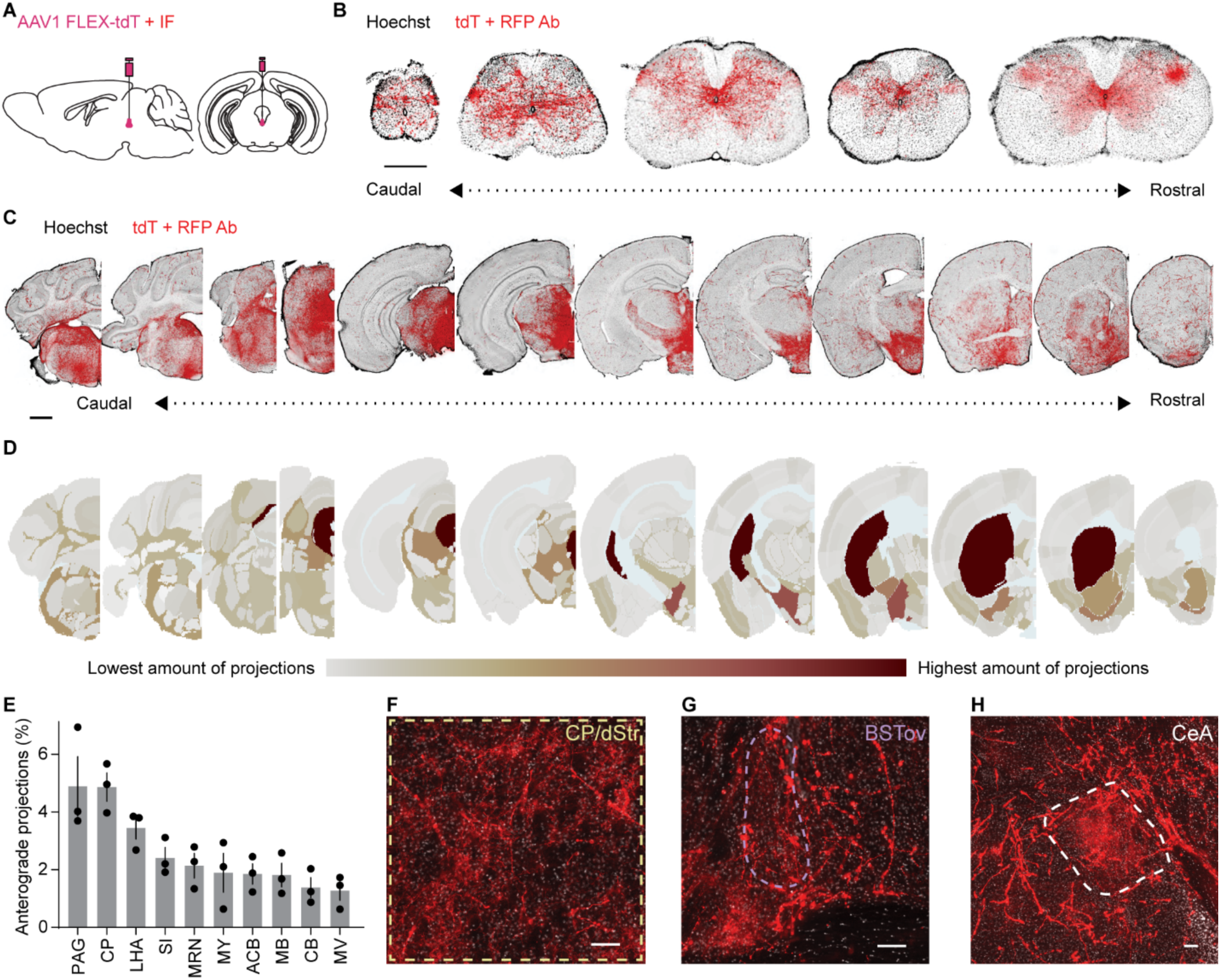
The CART^+^ EW projects to the spinal cord and multiple subcortical regions. (A) Schematic of AAV1 FLEX-tdT EW injection with immunoenhancement. (B) Coronal sections of CART^+^ EW projections to the spinal cord (coccygeal through cervical; Hoechst, black; immunoenhanced tdT, red; scale bar, 0.5 mm). (C) As in (B), but brain sections. Scale bar, 1 mm. (D) Projection intensity heat map across brain regions aligned to Allen Brain Atlas. Red, high expression; gray, low expression; light blue, undefined areas, white matter, ventricles. (E) Brain regions with the most CART^+^ EW projections (n = 3 mice), displayed as % of total quantified projections in the brain. (F-H) Immunoenhanced CART^+^ EW tdT in the (F) dorsal striatum/caudoputamen (dStr/CP), (G) oval nucleus of the bed nucleus of the stria terminalis (BSTov), and (H) the central nucleus of the amygdala (CeA). Scale bar, 100 µm. Error bars represent SEM. See also Figure S3.

The two brain regions with the greatest proportion of observed CART^+^ EW neuronal processes were the PAG and the caudoputamen (CP), also called the dorsal striatum (dStr) in mice (PAG: 4.89% ± 1.03%; CP/dStr: 4.87% ± 0.49%; n = 3 mice, Figures 3C to 3E). Given that the EW is located within the PAG, one possibility is that the numerous fibers we observe within the PAG are fibers of passage. However, within the dStr we observed wispy terminal fields, in addition to rarer varicosity-studded axonal fibers (Figure 3F). We observed similar terminal fields in the oval bed nucleus of the stria terminalis (BSTov) (Figure 3G) and in the central nucleus of the amygdala (CeA) (Figure 3H). While the CeA has many sizeable neuronal populations (Kim et al., 2017; McCullough et al., 2018), the BSTov has roughly three predominant populations of projection neurons (Daniel and Rainnie, 2016) and the dStr contains two large classes of striatal spiny projection neurons. Consequently, we focused on the dStr and BSTov to determine whether the CART^+^ EW affects the activity of neurons in these regions, and whether the modulation is consistent with a solely neuropeptide-mediated effect, or with glutamatergic effects.

### The CART^+^ EW is functionally peptidergic

To determine the nature of neurotransmitter release from CART^+^ EW neurons, we performed whole-cell current clamp on putative postsynaptic neurons in the dStr/CP and the BSTov, evoking vesicular release (Knobloch et al., 2012) from CART^+^ EW terminals through a single bout of optical activation of channelrhodopsin (ChR2) (10 ms, 10 mW, 30 Hz, 20 s) (Figure 4A). ChR2-driven stimulation of CART^+^ EW fibers more than doubled the firing rate in a subset of neurons in both the dStr/CP (7 of 16 SPNs) and the BSTov (3 of 11 cells) (Figures 4B, 4C, and S4A). In dStr spiny projection neurons (SPNs), SPNs with rheobase over 150 pA significantly increased excitability following CART^+^ EW stimulation (Figure 4D). To pharmacologically interrogate multipeptidergic release, we recorded from SPNs with rheobase over 150 pA and blocked intracellular signaling pathways downstream of GPCR activation. We observed that Gα_q_-signaling block using bath-applied phospholipase C inhibitor (U 73122) did not abolish SPN excitation induced by optogenetic stimulation of CART^+^ EW fibers (Figure 4D). However, increases in firing rate in SPNs were abolished by the intracellular block of Gα_s_-signaling with protein kinase A inhibitor (PKI) (Figure 4D). Increases in firing rate were also not found in the absence of ChR2 in CART^+^ EW neurons (Figure S4B).

**Figure 4:**
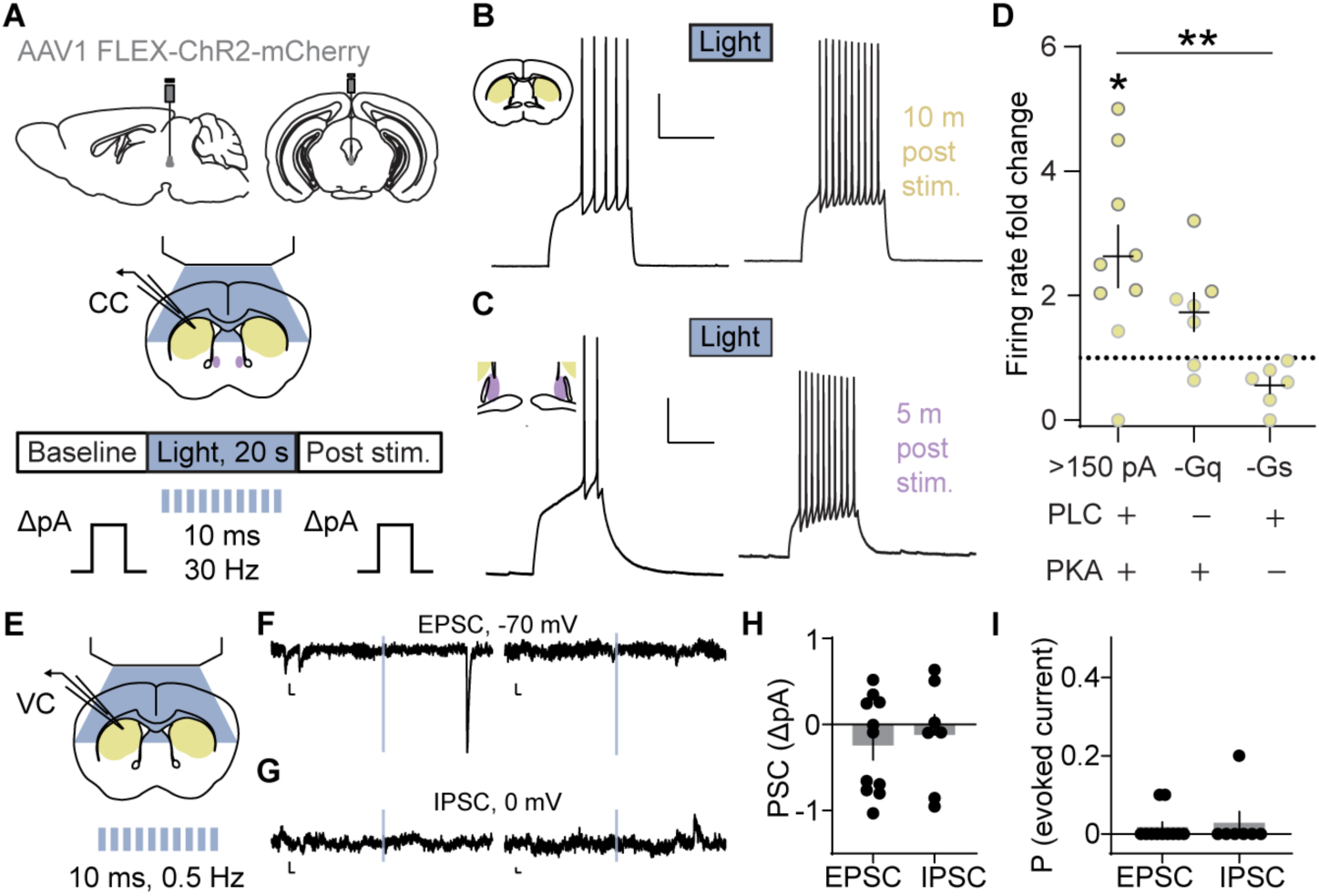
The CART^+^ EW is functionally peptidergic. (A) Schematic of AAV1 FLEX-ChR2-mCherry EW injection and whole-cell current clamp recordings of CART^+^ EW target regions (CP/dStr, yellow; BSTov, purple). An optical stimulation bout (20 s, 30 Hz) evokes vesicular release from ChR2-expressing CART^+^ EW terminals. (B) Following ChR2 stimulation, a striatal SPN fires more in response to a depolarizing current injection. Scale bars, 25 mV, 500 ms. (C) As in (B) but recorded from the BSTov. Scale bars, 20 mV, 500 ms. (D) SPNs with rheobase >150 pA generally increased their firing rate compared to baseline. *p < 0.05, paired t-test, t_(8)_ = 3.187, p = 0.0129, n = 9 neurons. Bath-applied PLC blocker (-Gq, n = 7 neurons) did not alter firing rate increases, but intracellular PKA block (-Gs, n = 6 neurons)) significantly decreased the light-evoked firing rate increase. **p < 0.01, one-way ANOVA with post hoc Dunnett’s test (F_(2,19)_ = 6.198; p = 0.0049; >150 pA vs. >150 pA -Gq, p = 0.2236; >150 pA vs. >150 pA -Gs, p = 0.0044). See also Figure S4. (E) Schematic of whole-cell voltage clamp recordings of CP/dStr with 0.5 Hz blue light applied in 10 ms pulses. (F) Example current traces from 2 SPNs as a 10 ms light pulse (blue) is applied while holding at -70 mV. Evoked EPSCs were not observed. Spontaneous EPSCs are observed. Scale bars, 5 pA, 20 ms. (G) As in (F), from SPNs held at 0 mV. Evoked IPSCs were not observed. Scale bars, 5 pA, 20 ms. (H) Summary of evoked current data, shown as the amplitude of postsynaptic current taken from the onset of the light pulse to 5 ms after the offset, compared to baseline (n = 11 neurons for EPSCs; 7 neurons for IPSCs). (I) Summary of evoked current data, shown as proportion of ten applied light pulses generating evoked EPSCs (n = 11 neurons) or IPSCs (n = 7 neurons). Error bars represent SEM.

To test whether the CART^+^ EW releases glutamate onto the downstream neurons that it excites, we pooled all neurons with a two-fold or greater firing rate increase following light-evoked vesicle release and examined membrane potential time-locked to optogenetic stimulation. In agreement with an absence of glutamatergic release, we observed no evidence of time-locked excitatory postsynaptic potentials (EPSPs) (Figures S4C and S4D). Finally, time-locked excitatory and inhibitory postsynaptic currents (EPSCs and IPSCs) were not observed in whole-cell voltage clamp recordings of SPNs in response to light stimulation (10 ms, 10 mW, 0.5 Hz, 5 pulses given in the first 10 s of a 24 s sweep, repeated twice) to evoke fast neurotransmitter release (Figure 4E and 4F). Taken together, our results suggest that the CART^+^ EW excites subpopulations of neurons in the BSTov and dStr and that the CART^+^ EW excitation of striatal projection neurons is dependent on intracellular Gα_s_-signaling cascades. The importance of Gα_s_-intracellular cascades for CART^+^ EW signaling aligns with the presence of numerous Gα_s_-linked peptides in the CART^+^ EW, including urocortin, PACAP, and CCK (Beinfeld et al., 2019; Dautzenberg et al., 2019; Fahrenkrug et al., 2019). Furthermore, we see no evidence that the CART^+^ EW is functionally glutamatergic. All anatomical and electrophysiological data support the thesis that the CART^+^ EW is a predominantly obligate peptidergic nucleus.

### The CART^+^ EW promotes fear responses

Next, we asked how the putative obligate peptidergic CART^+^ EW influences behavior. Numerous functions for the CART^+^ EW have been suggested, including consumption (Giardino et al., 2017), stress adaptation (Okere et al., 2010), and alertness (Li et al., 2018; Lovett-Barron et al., 2017). We chose to investigate the effect of the CART^+^ EW on fear responses, as both stress and alertness are linked to fear and many neuropeptides contained in the CART^+^ EW have been implicated in anxiety and fear (Bowers et al., 2012; Hammack and May, 2015; Stanek, 2006; Sztainberg and Chen, 2012). In addition, our data on terminal projections of the CART^+^ EW to the BSTov and CeA support a link to anxiety and fear responses. Additionally, the CART^+^ EW resides in the PAG, which responds to stressors, produces analgesia, and modulates defensive responses (George et al., 2019; Lefler et al., 2020; Silva and McNaughton, 2019).

Therefore, we tested whether CART^+^ EW activation or ablation alter fear- and anxiety-related behaviors. We conditionally expressed the Gα_s_-coupled, clozapine N-oxide (CNO)-activated rM3Ds for activation and diphtheria toxin for ablation of these neurons (Figures 5A, 5C, and 5D). Current-clamp recordings of rM3Ds^+^ CART^+^ EW neurons showed significant increases in firing rate following CNO application (Figures S5A and S5B) and diphtheria toxin expression routinely ablated >90% of EW neurons (Figure S5C). For controls, littermate mice of both sexes were infected with an inert control fluorophore, such as enhanced yellow fluorescent protein (EYFP) or tdT. Behavior was measured ≥3 weeks following AAV infection using a battery of behavioral assays to investigate fear and anxiety (Figure 5B). Mice were placed on an elevated plus maze, a behavioral assay for anxiety responses (Figure 5E). Mice with increased CART^+^ EW activity spent significantly less time on the open arms than littermate control AAV injected mice (Figure 5F), and conversely, mice with ablated CART^+^ EW neurons spent significantly more time on the open arms (Figure 5G).

**Figure 5:**
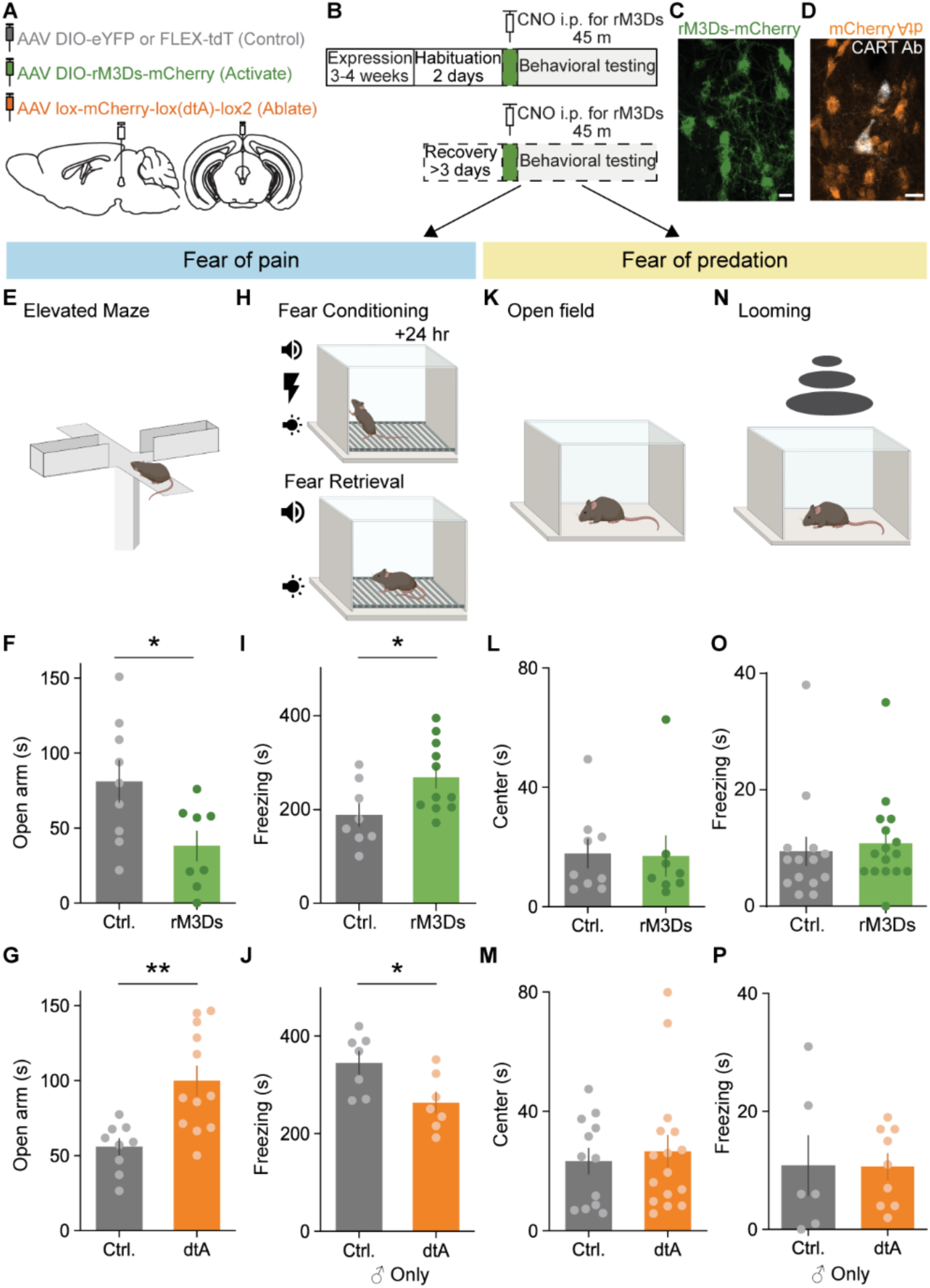
The CART^+^ EW promotes fear responses. (A) Schematic of injection of AAVs expressing Cre-dependent control fluorophore (gray), rM3Ds (green), or diphtheria toxin (dtA, orange). (B) Timeline of behavioral testing. (C) rM3Ds-mCherry (green) in CART^+^ EW. Scale bar, 20 µm. (D) Cre-off mCherry (orange) in the EW; CART immunofluorescence (white) confirms dtA ablates most CART^+^ EW neurons. Scale bar, 20 µm. (E) Elevated plus maze schematic. (F) CNO-injected rM3Ds-expressing mice spend less time on maze open arm than control mice. *p < 0.05, unpaired t-test, t_(15)_ = 2.481, p = 0.0255, n = 9 Ctrl. mice, 8 rM3Ds mice. (G) dtA mice spend more time on maze open arm than control mice. **p < 0.01, unpaired t-test, t_(19)_ = 3.575, p = 0.0020, n = 9 Ctrl. mice, 12 dtA mice. (H) Fear conditioning and retrieval schematic. (I) rM3Ds-expressing mice freeze more following conditioned cues than control AAV mice. *p < 0.05, unpaired t-test, t_(17)_ = 2.350, p = 0.0311, n = 8 Ctrl. mice, 11 rM3Ds mice. (J) Male dtA mice freeze less than male control mice. *p < 0.05, unpaired t-test, t_(12)_ = 2.570, p = 0.0245, n = 7 Ctrl. mice, 7 dtA mice. (K) Open field test schematic. (L) Time spent in center of the open field is similar for rM3Ds and control mice. Unpaired t-test, t_(15)_ = 0.1015, p = 0.9205, n = 9 Ctrl. mice, 8 rM3Ds mice. (M) As in (l), but dtA instead of rM3Ds. Unpaired t-test, t_(26)_ = 0.4580, p = 0.6507, n = 16 Ctrl. mice, 12 dtA mice. (N) Looming stimulus schematic. (O) As in (I), but for freezing response to looming stimulus. Unpaired t-test, t_(28)_ = 0.4422, p = 0.6618, n = 14 Ctrl. mice, 16 rM3Ds mice. (P) As in (O), but for dtA instead of rM3Ds, and for male mice only. Unpaired t-test, t_(13)_ = 0.0341, p = 0.9733, n = 6 Ctrl. mice, 9 dtA mice. Error bars represent SEM. See also Figure S5.

A separate cohort of mice underwent fear conditioning. On the first day, mice were exposed to four 30 s long paired light and auditory cues that ended with the onset of a 1 s shock. On the second day, mice were returned to the box and given the same cues another four times, with no paired shock (Figure 5H). Following fear conditioning, mice with CART^+^ EW activation froze significantly more (Figure 5I) and male mice (but not female mice) froze significantly less after ablation (Fig 5J). The observed changes in fear behavior are separable from changes in overall locomotor activity (Figures S5D to S5G); contextual fear measured immediately prior to the presentation of cue showed no difference between conditions (Figures S5H and S5I).

Interestingly, we did not observe promotion of fear by the CART^+^ EW in all behavioral assays. For example, perturbations of the CART^+^ EW did not alter the amount of time spent in the arena center during an open field test (Figures 5K to 5M), nor did it alter behavior in other anxiety tests such as a light-dark box test and a novel arena suppressed feeding test (Figures S5J and S5K), or in response to a looming stimulus (Yilmaz and Meister, 2013) (Figures 5N to 5P). One noticeable difference between the behavioral assays that the CART^+^ EW influenced and those it did not is the degree to which the fear- or anxiety-inducing stimuli reflect the risk of pain, rather than the threat of predation. For example, there is nothing inherently painful about an open-field test, a light-dark box test, or a novel arena suppressed feeding test. Instead, the combination of bright lights and open, novel spaces are anxiogenic as the risk of threats including predators is increased (Calhoon and Tye, 2015). Similarly, the looming stimulus is designed to evoke a predator approaching the animal (Yilmaz and Meister, 2013). The conditioned fear test, on the other hand, clearly measures fear responses to stimuli associated with pain, while the elevated maze combines the animals’ fear of open spaces with their fear of heights and the risk of a potentially painful fall (Walf and Frye, 2007). Recently, it has been suggested that fear of pain is mediated by neural pathways that are separable from those mediating fear of predation or social interaction fear (Gross and Canteras, 2012; Silva et al., 2013). One parsimonious explanation for the observed CART^+^ EW-induced behavioral responses is that these neurons selectively promote responses related to a fear of pain, rather than a fear of predation.

### The CART^+^ EW receives inputs from brain regions related to motor control and threat responses

In order to better understand the circuitry giving rise to behavioral control of fear responses by the CART^+^ EW, we characterized inputs to the CART^+^ EW through the whole central nervous system using monosynaptic retrograde mapping (Figure 6A). GFP-expressing helper virus colocalized with tdT- expressing pseudotyped rabies virus strain CVS-N2c (Reardon et al., 2016) in the EW (Figure S6A). Previously, the ventral hippocampus, medial septum, dorsal raphe, locus coeruleus (Li et al., 2018), and the ventral tegmental area (Ryabinin et al., 2013) had been claimed to project to the genetically defined neuropeptidergic Edinger-Westphal. We did not find consistent evidence (≥2 neurons in each mouse) of projections from any of these areas despite observing sparse labeling in nearby areas, e.g., the dorsal subiculum and lateral septum (Figure S6B to S6E, n = 3 mice, 190 to 233 neurons per mouse). Notably, despite a prior claim that glutamatergic neurons in the ventral hippocampus provide monosynaptic excitatory input to the EW (Li et al., 2018), we found zero retrogradely labeled neurons in this area (Figures S6B and S6E). We did, however, find robust and consistent evidence of CART^+^ EW inputs in numerous cortical and subcortical regions of the brain (Figure 6B).

**Figure 6:**
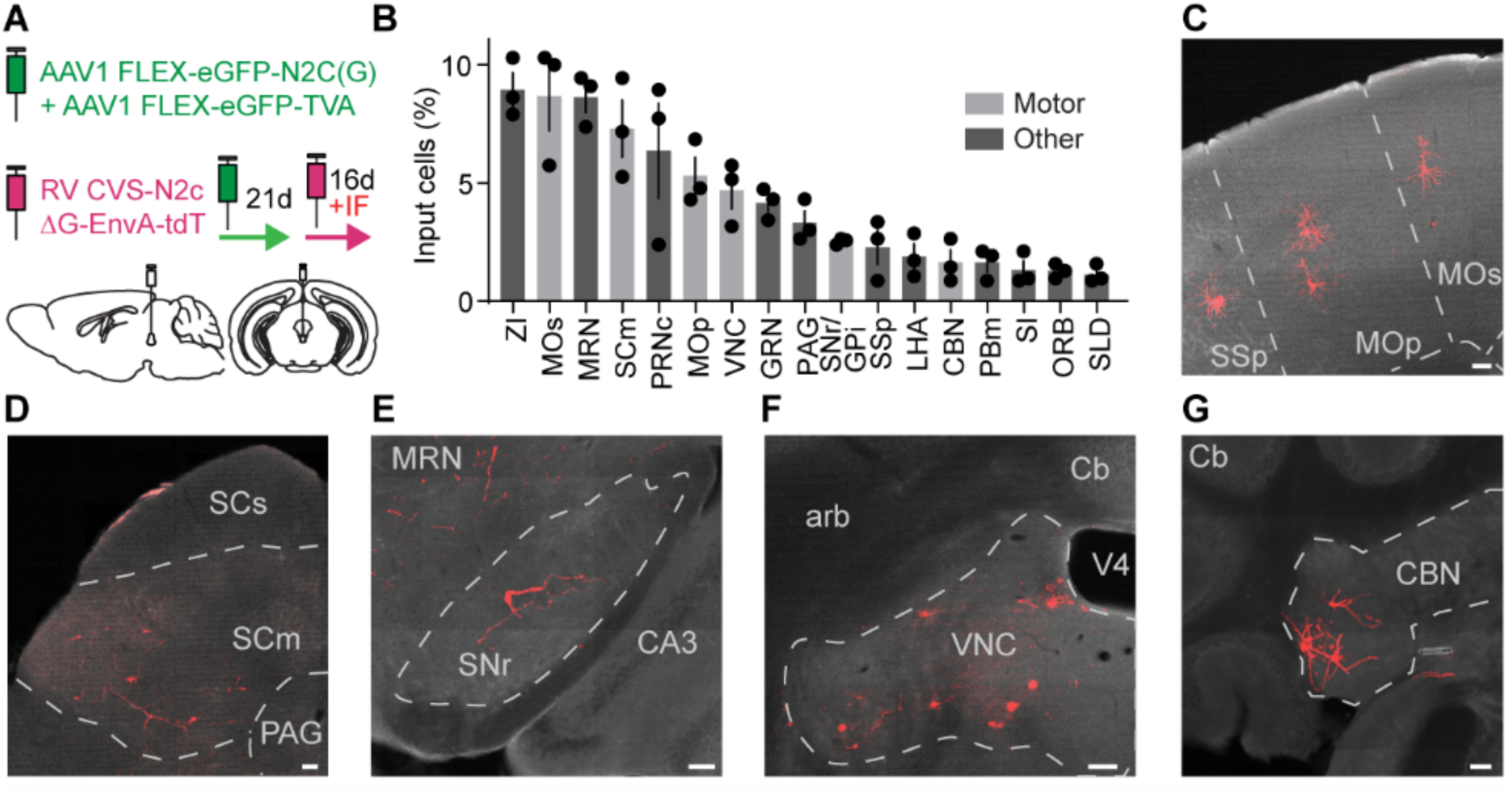
The CART^+^ EW receives inputs from brain regions related to motor control and threat responses. (A) Schematic of monosynaptic retrograde tracing from CART^+^ EW. tdT fluorescence is immunoenhanced. (B) Retrogradely labeled tdT^+^ neurons by brain region, as percentage of all retrogradely labeled tdT^+^ neurons (n = 3 mice, 190 to 233 neurons per mouse). Motor control brain regions are in gray. (C) Image of retrogradely labeled neurons (red) in primary motor cortex (MOp), secondary motor cortex (MOs), and primary somatosensory cortex (SSp). Scale bar, 100 µm. (D-G), as in (C), but retrogradely labeled neurons in (D) motor related areas of superior colliculus (SCm), (E) substantia nigra pars reticulata (SNr), (F) vestibular nuclei (VNC), and (G) cerebellar nuclei (CBN). Error bars represent SEM. See also Figure S6.

Genetically-targeted retrograde tracing does not provide information on synaptic input strength (Ugolini, 2011). Therefore, we examined whether any of the 17 brain regions consistently found to provide input to the CART^+^ EW shared a common function, rather than focusing solely on the regions where the highest number of neurons were found. We found monosynaptic CART^+^ EW inputs from 6 brain regions directly related to motor control (Figure 6E): pyramidal neurons in layer V of the primary and secondary motor cortices (MOp, MOs) (Figure 6F), motor-related superior colliculus (SCm) (Figure 6G), substantia nigra pars reticulata (SNr) (Figure 6H), vestibular nuclei (VNC) (Figure 6I), and cerebellar nuclei (Figure 6J). This enrichment of input nuclei involved in motor control is similar to that observed for the input nuclei of the substantia nigra pars compacta (SNc), a neuromodulatory nucleus considered vital for motor control. 6 of the 17 brain regions with the greatest number of labeled cells found upstream of the SNc using monosynaptic retrograde labeling are intimately tied to motor control (Watabe-Uchida et al., 2012).

In addition to receiving direct inputs from cortical and subcortical regions that transmit information about motor commands or outcomes, we found that the CART^+^ EW also received projections from multifunctional regions involved in responses to pain or threat, including the somatosensory cortex (Iwamoto et al., 2021), lateral hypothalamus (Chen et al., 2020; Siemian et al., 2021), zona incerta (Chou et al., 2018; Zhou et al., 2018), and periaqueductal gray (George et al., 2019; Lefler et al., 2020) (Figure 6B).

### CART^+^ EW neurons respond to loss of motor control

To determine the relationship between CART^+^ EW neuronal activity and movement, we expressed GCaMP6s and measured *in vivo* CART^+^ EW activity with angled mirror tipped photometry fibers to avoid the cerebral aqueduct (Figures 7A, 7B and S7A). Under blue light (470 nm) excitation of GCaMP6s, we observed that gentle tail restraint elicited calcium transients that were time-locked with the restraint and reproducible across tail restraint trials and different mice (Figure 7C). To control for the possibility of motion artifact induced transients, we performed separate trials using light (405 nm) that excites near the isosbestic point. No transients were observed under isosbestic illumination (Figures S7B to S7E), suggesting that the transients we observed under blue light excitation are due to changes in intracellular calcium concentration in the CART^+^ EW. Similar fluorescence signals were observed in response to tail suspension (Figures 7D and S7C).

**Figure 7:**
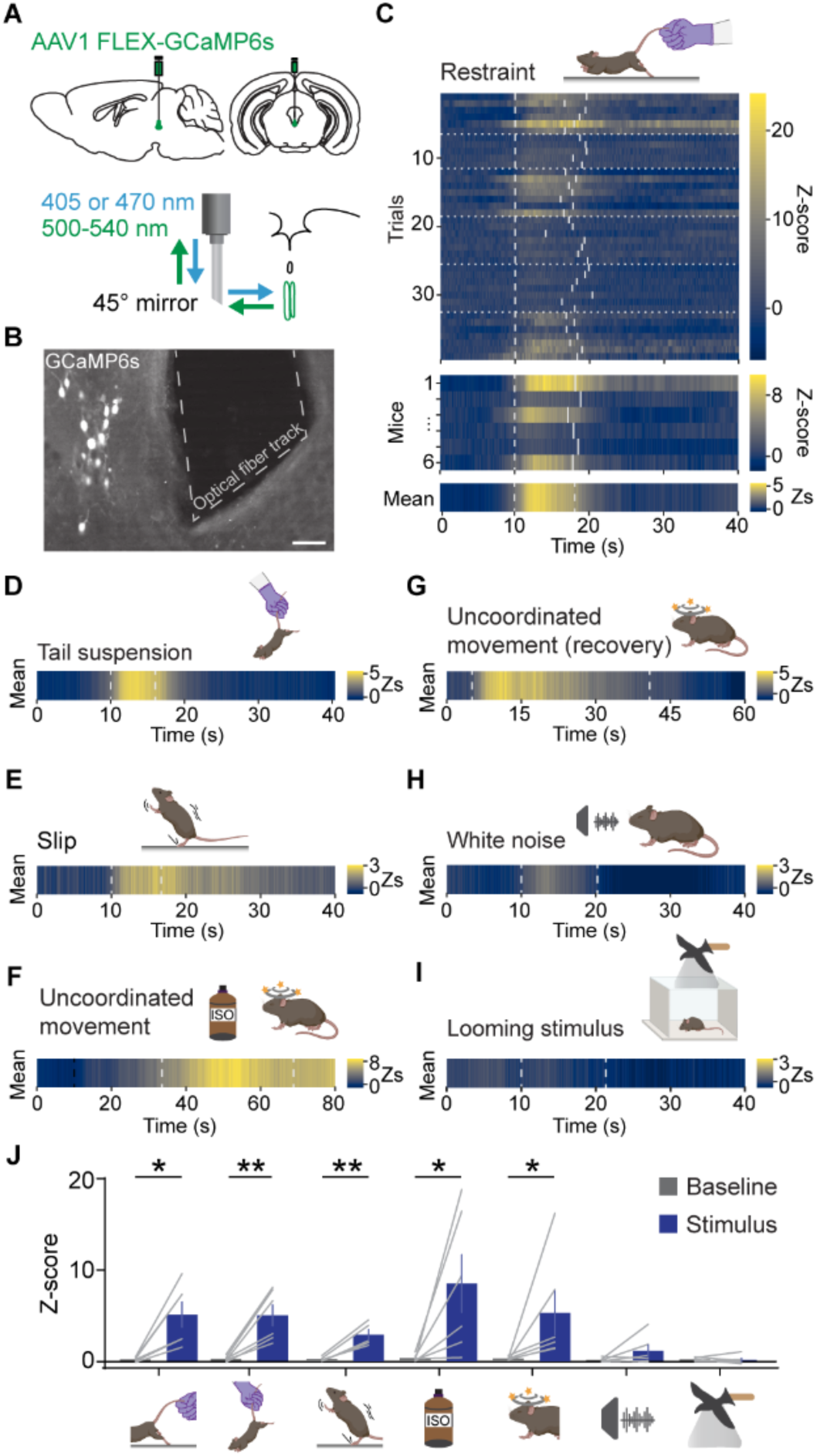
CART^+^ EW neurons respond to loss of motor control. (A) Schematic of CART^+^ EW fiber photometry for *in vivo* calcium recordings. A 45° angled mirror tip fiber is used to avoid occluding the cerebral aqueduct dorsal to the CART^+^ EW. 405 and 470 nm light measures motion artifact induced fluorescence and calcium concentration dependent fluorescence, respectively. (B) GCaMP6s expression and 45° angled mirror tip fiber placement. Scale bar, 50 µm. (C) Time-locked fluorescence signals recorded from the CART^+^ EW in response to mild tail restraint. Heatmap of z-scores is from blue (little to no calcium signal) to yellow (high calcium signal). Top, cartoon of tail restraint. Below, each trial (n = 5 to 7) from each mouse (n = 6). The start and end of each manual tail restraint are shown by dashed gray lines. Data from different mice are separated by dotted gray lines. Beneath are fluorescence signals for each mouse, averaged across all trials. The stimulus time points are shown by dashed gray lines marking the average start and stop point of all trials for a given animal. Bottom, the fluorescence signal (Zs, or Z-score) averaged across all mice, with the stimulus time points shown as a dashed gray line at the average start and stop point across all animals. (D) As in the bottom panel of (C), but in response to tail suspension (n = 6 mice, 6-9 trials each). (E) As in the bottom panel of (C), but in response to slipping following placement in an arena with a thin layer of corn oil (n = 5 mice, 4 slipping bouts each) (F) As in the bottom panel of (C), but in response to anesthetic isoflurane exposure (n = 6 mice, 3-4 trials each). Exposure to isoflurane (dashed black line) does not elicit time-locked fluorescence signals. Uncoordinated movement following isoflurane exposure (dashed gray line) coincides with GCaMP6s fluorescence transients. (G) As in the bottom panel of (C), but in response to uncoordinated movement during recovery from anesthesia (n = 6 mice, 3-4 trials). Average start and stop times of uncoordinated movement are marked by dashed gray lines. (H-I) As in the bottom panel of (C), but in response to loud white noise (H) (n = 6 mice, 6 to 8 trials each) or looming stimulus (I) (n = 6 mice, 6 to 7 trials each). Dashed gray lines mark the onset and offset of the noise. The Zs heat map is set to match the heat map used in the slipping data (∼-1 z to 3 z), as this was the stimulus that reproducibly activated the CART^+^ EW with the smallest fluorescence changes. (J) Peak z-score of stimulus-driven fluorescence changes versus baseline for, from left to right, restraint (*p<0.05, t(5) = 3.80, p=0.0126), tail suspension (**p<0.01, t(5) = 4.469, p=0.0066), slip (**p<0.01, t(4) = 5.83, p=0.0043), isoflurane-induced uncoordinated movement (*p<0.05, t(5) = 2.64, p=0.0459), uncoordinated movement during recovery from anesthesia (*p<0.05, p=0.0313), white noise (t(5) = 1.526, p=0.1876), and looming stimulus (t(5) = 0.2596, p=0.8055). Paired t-test used for all comparisons except uncoordinated movement during recovery from anesthesia (Wilcoxon matched-pairs signed rank test), based on a Shapiro-Wilk test of normality. Error bars represent SEM.

We also observed that CART^+^ EW calcium transients were sporadic while the mouse moved freely in its home cage or an open field (0.017 Hz ± 0.008 Hz, n = 6 mice, z-score ≥ 3). We noticed that transients coincided with brief losses of balance or slipping by the animal (29.2% ± 14.3% of observed transients, n = 5 mice, 17 of 50 observed transients). We confirmed this by placing mice in an arena with a thin film of corn oil: we observed time-locked fluorescence signals when mice slipped (Figure 7E).

Interestingly, induction of CART^+^ EW activity has previously been observed through increases in c-fos following anesthetic exposure (Gaszner et al., 2004). We found that CART^+^ EW neurons activate during the transient motor dysfunction that occurs following anesthetic exposure (Figure 7F). Furthermore, CART^+^ EW fluorescence signal also increased as the mice moved in an uncoordinated fashion while recovering from anesthesia (Figure 7G). Thus, we observed increases in CART^+^ EW activity following multiple stimuli that induce a loss of motor control, whether passively (slipping on oil), actively (restraint or suspension), or chemically (anesthetic exposure) (Figure 7J). Importantly, other stressful stimuli that did not alter the motor control of the animal, such as a loud white noise (Figure 7H) or looming stimulus (Figure 7I), did not reproducibly elicit time-locked fluorescence responses (Figure 7J). The bouts of motor control loss that we observe activating the CART^+^ EW could also increase the threat of pain for the animal, aligning with the promotion of fear of pain we observe upon EW activation (Figure 5).

### The CART^+^ EW is pain-responsive and analgesic

In addition to the responses to motor control loss and increased threat of pain we have shown here (Figure 7), the CART^+^ EW has previously been suggested to respond to acute pain (Innis and Aghajanian, 1986; Kozicz et al., 2001). CART^+^ EW neurons receive monosynaptic input from brain regions implicated in pain processing including primary somatosensory cortex, zona incerta, and the periaqueductal gray (Figure 6), and we localized CART^+^ EW projections to pain processing regions including the central amygdala (Han et al., 2015; Wilson et al., 2019) and laminae III, IV, and X of the spinal cord (Figure 3 and S3). Finally, the PAG, which encompasses the CART^+^ EW, has been shown to mediate both opioidergic and non-opioidergic analgesia (Keay and Bandler, 2015). Therefore, in addition to motor control loss-associated pain threat, we tested the responsiveness of the CART^+^ EW to brief painful stimuli.

Using fiber photometry as before, we found that the CART^+^ EW activates following electric shocks but not to conditioned fear cues (Figures 8A and S8A to S8C). In response to a von Frey test of paw withdrawal to mechanical stimuli of different force levels, activation of the CART^+^ EW significantly increases paw withdrawal threshold, suggesting that these neurons promote analgesia (Figure 8B). This suggests that the fear of pain induced by the EW (Figure 5) is not a consequence of increased pain sensitivity. We found that despite its numerous neuropeptides, most neurons of the CART^+^ EW lack the opioid-encoding mRNAs for proenkephalin (*Penk*, 4.44% ± 1.93%, n = 3 mice, 385 *Cart*^+^ neurons), prodynorphin (*Pdyn*, 3.22% ± 1.55%, n = 3 mice, 385 *Cart*^+^ neurons), and proopiomelanocortin (*Pomc*, 1.40% ± 0.65%, n = 3 mice, 636 *Cart*^+^ neurons) (Figures 8D, 8E, and S8D to S8F). Thus, the analgesia produced by EW activation does not stem from direct release of opioids onto downstream cellular targets.

**Figure 8:**
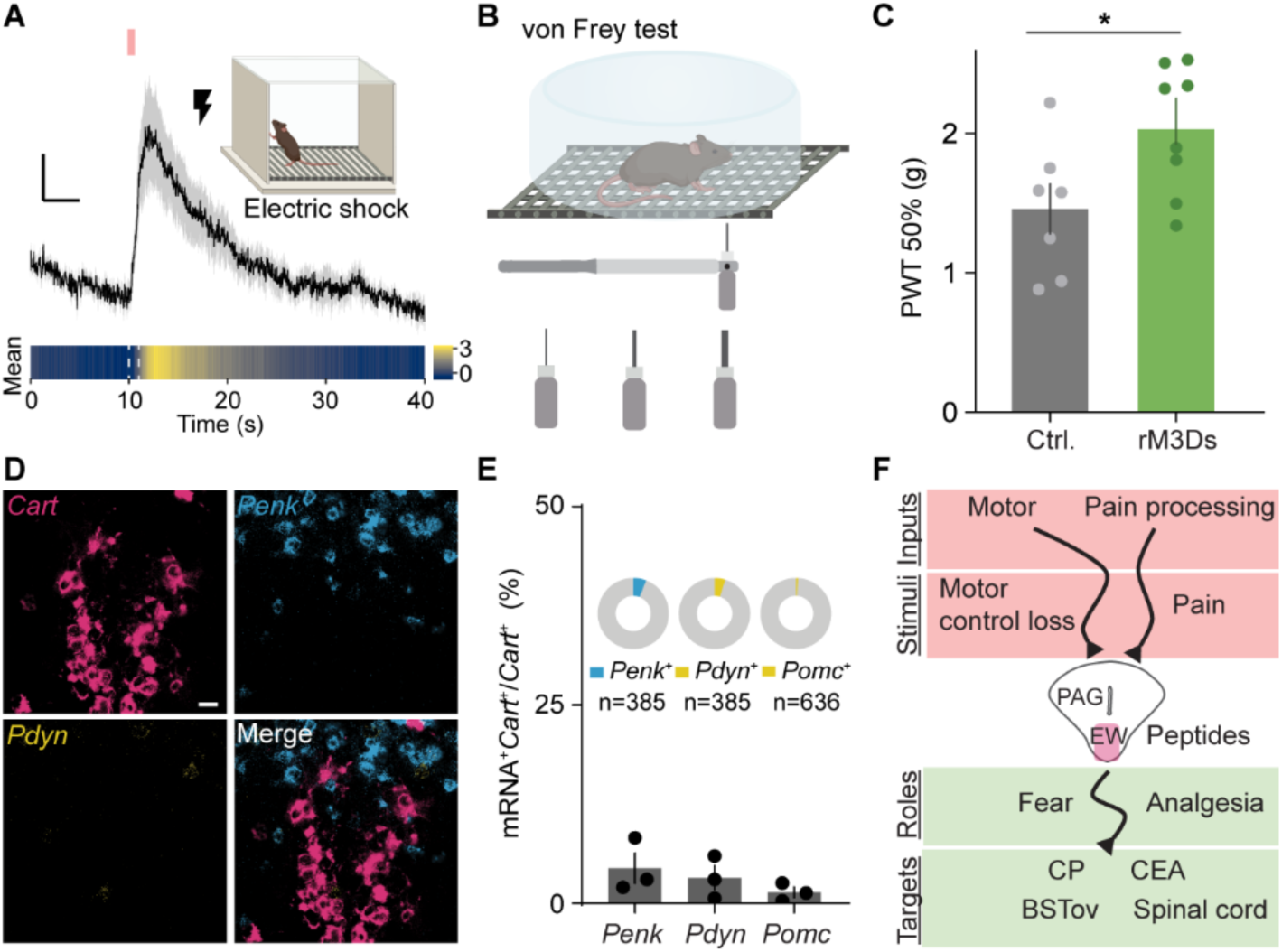
The CART^+^ EW is pain-responsive and analgesic. (A) Top, CART^+^ EW GCaMP6s fluorescence signal time-locked to painful electric shock (pink). Mean from 6 mice, 7 trials per mouse. Scale bar, 1 z, 5 s. Bottom, the same data as a heatmap, with electric shock marked by gray dashed line. (B) Von Frey paw withdrawal test cartoon. (C) rM3Ds mice show decreased mechanical sensitivity compared to controls (*p < 0.05, Mann-Whitney test, p = 0.0401 n = 7 Ctrl. mice, 8 rM3Ds mice). (D) FISH of EW *Cart* (red) against *Pdyn* (yellow) and *Penk* (blue). Scale bar, 20 µm. (E) EW *Cart* does not colocalize with opioid-encoding mRNAs (*Penk*, 4.44% ± 1.93%; *Pdyn*, 3.22% ± 1.55%, *Pomc*, 1.40% ± 0.65%, n = 3 mice, inset pie graphs show pooled *Cart*^+^ count). (F) An anatomical-, circuit-, and function-based model for the EW as a ventromedial column of the PAG. Gray shading in (A) and error bars represent SEM.

Summarizing our data, we propose that the CART^+^ EW responds to stimuli that are painful or increase the threat of pain through a loss of motor control. It then signals in a predominantly obligate peptidergic fashion to numerous downstream targets throughout the central nervous system, enhancing fear responses to pain or increased risk of pain (Figure 8F).

## Discussion

We hypothesize that neuronal populations in the brain including the CART^+^ EW could be obligately peptidergic, i.e., incapable of fast neurotransmission. Proving the absence of something is intrinsically difficult, especially given the existence of non-canonical purinergic and gaseous transmitters, and further research will be needed to formally define any neuronal population as obligately peptidergic. For example, prior studies of Trh^+^, Oxt^+^, and Avp^+^ neurons of the PVN, have potentially ruled out these populations as being obligately peptidergic. Robust monosynaptic EPSCs have been recorded in agouti-related peptide (AgRP) neurons of the arcuate hypothalamus upon optogenetic stimulation of Trh^+^ PVN terminals, but glutamatergic transmission from Oxt^+^ or Avp^+^ neurons was not observed (Krashes et al., 2014). Previously, we did not observe glutamatergic transmission from Oxt^+^ terminals in the ventral tegmental area (Xiao et al., 2018). Following ChR2 expression in the PVN, fibers that projected to the dorsal motor nucleus of the vagus included Oxt^+^ fibers and released glutamate (Piñol et al., 2014). Similarly, expression of ChR2 in a mixed population of Oxt^+^ and oxytocin receptor^+^ neurons of the PVN could elicit light-evoked action potentials and EPSCs in a subpopulation of oxytocin receptor^+^ neurons of the parabrachial nucleus (Ryan et al., 2017). Others have found that the Oxt^+^ PVN is glutamatergic, based on Vglut2 immunolabeling in Oxt^+^ PVN fibers and glutamate-dependent modulation of disynaptic IPSCs following light-evoked excitation of these fibers (Grinevich and Ludwig, 2021; Hasan et al., 2019; Knobloch et al., 2012). Finally, the *Avp*^+^ PVN has recently been found to colocalize with *Vglut2* based on FISH (Zhang et al., 2020). However, all our FISH, immunofluorescence, electron microscopy, and electrophysiology data suggest that the CART^+^ EW may be an obligate peptidergic population.

One piece of evidence against our hypothesis is a prior report that evoked EPSCs on parvalbumin interneurons of the medial prefrontal cortex (mPFC) with optogenetic stimulation of ChR2^+^ fibers following delivery of ChR2 to the EW of a CCK-Cre transgenic mouse line (Li et al., 2018). We cannot preclude that the small proportion of CART^+^ neurons we found containing mRNA for a vesicular glutamate transporter (∼11.5%) project to the mPFC, although we never observed robust projections to this brain region. Another possible explanation is that the use of AAVs with a CCK-Cre mouse line may not selectively target the CART^+^ EW, even though CART and CCK colocalize in these neurons. For example, AAV injection in the midbrain of a CCK-Cre mouse line drove expression of ChR2 in sleep-promoting glutamatergic CCK^+^ neurons in the perioculomotor region (Zhang et al., 2019). Additionally, characterization of a CCK-Cre mouse line (Taniguchi et al., 2011) shows robust expression in the mediodorsal nucleus of the thalamus. Mediodorsal thalamic neurons supply robust glutamatergic inputs to parvalbumin interneurons in medial prefrontal cortex (Delevich et al., 2015), and could underlie the putatively-EW derived EPSCs previously recorded from the mPFC (Li et al., 2018).

The CART^+^ EW has a history of being mischaracterized. The anatomical area now shown to contain the CART^+^ EW was incorrectly defined in the 19^th^ and much of the 20^th^ century as the site of the cholinergic Edinger-Westphal oculomotor nucleus (Kozicz et al., 2011). As a result, despite residing within the PAG, the CART^+^ EW has typically been considered distinct (Keay and Bandler, 2015). Both the CART^+^ EW and the PAG are phylogenetically ancient, with strong conservation through mammals (Kozicz et al., 2011; Silva and McNaughton, 2019) and proposed homologous structures found in teleost fish (Lovett-Barron et al., 2017; Silva and McNaughton, 2019). The PAG is known to mediate defensive behaviors in response to threats and pain, producing fear responses and analgesia (George et al., 2019; Gross and Canteras, 2012; Keay and Bandler, 2015; Lefler et al., 2020; Silva and McNaughton, 2019), and we show here that the CART^+^ EW responds to loss of motor control, which could threaten pain, as well as pain itself, and that this peptidergic nucleus promotes fear responses and analgesia. The PAG canonically comprises four columns: dorsomedial, dorsolateral, lateral, and ventrolateral (George et al., 2019; Silva and McNaughton, 2019), with different columns potentially responding preferentially to specific classes of threat (Gross and Canteras, 2012; Keay and Bandler, 2015). The phylogenetic and anatomical similarities between the CART^+^ EW and the PAG, as well as our *in vivo* fiber photometry and behavioral assay results, yield a model in which the CART^+^ EW behaves as a ventromedial column of the PAG. Additionally, the results of this study clarify opposing characterizations of CART^+^ EW function, advance our understanding of neuromodulatory mechanisms underlying fear behavior, and support the classification of the CART^+^ EW as an obligate peptidergic nucleus.

Numerous questions surrounding peptidergic signaling remain, including details surrounding the genesis and maturation of large vesicles (Ailion et al., 2014; Kim et al., 2006), to what extent peptides are colocalized or segregated within individual vesicles or release sites (Merighi, 2018), the mechanisms underpinning peptide vesicle release (Chang et al., 2021; Ding et al., 2019; Moro et al., 2021; Persoon et al., 2019; Puntman et al., 2021), and the modulatory capabilities of neuronal processing in the absence of faster-acting neurotransmitters (Nusbaum and Blitz, 2012). Our characterization of a putatively obligate peptidergic population in the mammalian midbrain, together with associated ethologically relevant behaviors, provides an effective system for investigation of neuronal peptide transmission.

## Acknowledgments

We thank the Northwestern University Biological Imaging Facility, as well as Dr. Tiffany Schmidt and Dr. Reza Vafabakhsh for confocal microscope access, the Northwestern University BioCryo Facility of the NUANCE Center for TEM sample preparation and microscope use, Dr. Martha Vitaterna for loaning behavioral apparatuses (elevated mazes, open field), Dr. Gregory Scherrer for a gift of biotinylated IB4, Lindsey Butler for mouse colony management, and Jordan Nasenbeny and Sara Boyle for technical assistance in early stages of the study. This work was supported by NIH R01MH117111, NIH R01NS107539, NSF CAREER Award 1846234, and the following awards: Beckman Young Investigator, Rita Allen Foundation Scholar, Searle Scholar, William and Bernice E. Bumpus Young Innovator, NARSAD Young Investigator/P&S Fund (to Y.K.). M.F.P was supported by the Arnold O. Beckman Postdoctoral Fellowship and NIH T32AG20506. S.N.F. was supported by NIH T32MH067564. V.D. was supported by an American Heart Association predoctoral fellowship (19PRE34380056) and by NIH T32GM15538.

## Author contributions

M.F.P. and Y.K.: Conceptualization, Methodology, Software, Validation, Investigation, Resources, Data Curation, Writing – original draft, Writing – review & editing, Visualization. S.N.F., D.B., and V.D.: Methodology, Software, Validation, Investigation, Resources, Data Curation, Writing – review & editing, Visualization. Y.K.: Supervision, Project administration, Funding acquisition.

## METHODS

### Mouse lines and husbandry

Mice used in all experiments except fluorescence *in situ* hybridization were heterozygous, formed by crossing B6;129S-Cartpttm1.1(cre)Hze/J^+/-^ (CART-Cre, #028533, The Jackson Laboratory) (Daigle et al., 2018) mice to each other, or, more commonly, crossing CART-Cre^+/+^ mice to wild-type C57BL/6J mice (Charles River, Wilmington, MA). Wild-type C57BL/6J mice were used for fluorescence *in situ* hybridization. After weaning, experimental mice were put in single-sex housing with ad libitum food and water. Mouse were generally maintained on a 12h:12h light-dark cycle, with a subset of mice maintained on a 12h:12h reverse light-dark cycle following surgery prior to behavioral experiments. Male and female mice were used for all experiments, and all experiments were performed on adult (>P40) mice. Littermates of the same sex were randomly assigned to experimental groups, when applicable. All mouse handling, surgeries, and behavioral experiments were performed according to protocols approved by the Northwestern University Animal Care and Use Committee.

### Single-cell RNAseq database analysis

The dataset ’l6_r2_cns_neurons.loom’ was downloaded from mousebrain.org (Zeisel et al., 2018). Neuronal types were taken from the clusters defined within the scRNAseq database. A custom MATLAB (MathWorks) script tabulated the existence of user-defined transcripts in each cell, where any evidence of a transcript in a cell was sufficient to positively-identify a cell as containing that transcript. If a cell was positively identified as containing a transcript that was a marker for any population (i.e., glutamatergic, GABAergic, or monoaminergic), it was considered positively identified within that population. The proportion of cells in each neuronal type that were positively identified for a given neurotransmitter-defined population were calculated, and neuronal types were then plotted in three dimensions based on these proportions.

### Fluorescence in situ hybridization

Wild-type C57BL/6J mice were deeply anesthetized prior to decapitation and brain extraction. Brains were rapidly frozen in Tissue-Tek O.C.T. Compound (VWR) using a slurry of dry ice and ethanol and then transferred to -80 °C overnight. 20 μm thick brain slices for fluorescence in situ hybridization were cut from fresh frozen brains at -15 °C to -25 °C using a Leica CM1850 cryostat (Leica Biosystems). Slices were mounted on Superfrost Plus microscope slides (Fisher Scientific) and processed and labeled with fluorescence in situ hybridization probes according to the manufacturer (ACDBio) instructions. Probes used included Cartpt-C1, Adcyap1-C1, Penk-C1, Nmb-C1, Slc17a6-C2, Slc17a7-C2, Cck-C2, Pdyn-C2, Pomc-C2, Cartpt-C3, Tac1-C3, and Slc32a1-C3. Labeled slices were covered with Prolong Gold Antifade Mountant with DAPI (ThermoFisher Scientific) and coverslipped.

FISH data were collected and analyzed similarly to previous descriptions (Xiao et al., 2017, 2018). Stacks were taken at a 0.5 µm interval on a Zeiss 880 or Leica SP8 confocal microscope at 40x, with imaging for DAPI, AlexaFluor 488, Atto 550, and Atto 647. Five consecutive z-plane images were merged for analysis, and cells were found based on DAPI signal and a watershed algorithm on fluorescent signal. *Cartpt* transcript was frequently so abundant that individual puncta could not be observed at this or higher magnification levels; a conservative checkerboard counting system was used (Xiao et al., 2018), assuming that the fluorescence signal from puncta were approximately 2 pixels (1.248 µm) in diameter.

### Intracranial injections and implants

Viral vectors were stored at -80 °C prior to use and were backfilled into Wiretrol II pipettes (Drummond Scientific Company) pulled on a P-1000 Flaming/Brown micropipette puller (Sutter Instruments). Mice were anesthetized with vaporized isoflurane and positioned on a stereotactic apparatus (Kopf Instruments) so the skull was lying flat with the vertical positions of lambda and bregma within 0.1 mm of each other, and the pipette was placed at the midline (±0.0 mm M/L), 0.7 mm rostral to lambda (+0.7 mm A/P), and, generally, both 3.5 and 3.1 mm ventral to the pial surface (-3.5 and -3.1 D/V). Virus was injected using a microsyringe pump controller (World Precision Instruments); >2 minutes elapsed before the pipette was moved from the ventral injection site to the more dorsal injection site, and >5 minutes elapsed before the pipette was slowly retracted fully from the brain following the second injection. Mice were given ketofen (5 mg/kg) for analgesia and recovered on a heating pad before being returned to their home cage.

For anterograde fluorescent projection mapping, 50-100 nl of AAV1-CAG-FLEX-tdTomato-WPRE (1.9 x 10^13^ gc/ml, University of Pennsylvania Vector Core) was injected at -3.5 and -3.1 D/V at a rate of 50 nl/min. For APEX-mediated electron microscopy, 200 nl of AAV1-EF1α-DIO-LCK-APEX2-P2A-EGFP (University of North Carolina Vector Core, 5.0 x 10^12^ gc/ml) was injected at -3.3 D/V at a rate of 100 nl/min. For light-induced excitation, 200-250 nl of AAV1-CBA-FLEX-ChR2-mCherry (Atasoy et al., 2008) (5 to 6 x 10^12^ gc/ml, University of Pennsylvania Vector Core or Vigene) was injected at -3.5 and -3.1 D/V at a rate of 100 nl/min. For pharmacogenetics experiments, 300 nl of AAV1-CBA-DIO-rM3Ds-mCherry- WPRE (Wu et al., 2021a) (Vigene, 1.08 to 4.3 x 10^13^ gc/ml), was injected at -3.5 and -3.1 D/V at a rate of 100 nl/min.

For retrograde rabies tracing, AAV1-CAG-Flex-H2B-eGFP-N2c(G) (1.43 x 10^12^ gc/ml, Columbia Vector Core) and AAV1-EF1α-FLEX-GT (6.14 x 10^11^ gc/ml, Salk Institute Viral Vector Core) were injected into adult mice in a single 150-200 nl injection at -3.4 D/V at a rate of 100 nl/min. After three weeks, rabies virus CVS-N2cΔG tdTomato EnvA (Reardon et al., 2016) (Columbia Vector Core) was injected at -3.6, - 3.3, and -3.0 D/V with 400 nl released at each site at a rate of 100 nl/min.

For genetically targeted ablation, 400 nl of AAV1-EF1α-Lox-mCherry-lox(dtA)-lox2 (Wu et al., 2014) (University of North Carolina Vector Core, 5.2 x 10^12^ gc/ml) was injected at -3.5 and -3.1 D/V at a rate of 100 nl/min. Control behavioral animals were injected with AAV1-CAG-FLEX-tdTomato-WPRE or AAV9-EF1α-DIO-eYFP-WPRE in a titer and volume matched manner (Addgene or University of Pennsylvania Vector Core). For fiber photometry, 200-250 nl of AAV1-CAG-FLEX-GCaMP6s-WPRE (Chen et al., 2013) (University of Pennsylvania Vector Core, 4 x 10^12^ to 1 x 10^13^ gc/ml) was injected at - 3.5 and -3.1 D/V at a rate of 100 nl/min.

AAV1-CBA-FLEX-ChR2-mCherry was packaged from AAV-FLEX-rev-ChR2(H134R)-mCherry plasmid, which was a gift from Scott Sternson (Addgene plasmid #18916). AAV1-CBA-DIO-rM3Ds-mCherry-WPRE was packaged from plasmid pAAV-hSyn-DIO-rM3D(Gs)-mCherry, which was a gift from Bryan Roth (Addgene plasmid #50458). AAV1-CAG-FLEX-GCaMP6s-WPRE was packaged from plasmid pAAV.CAG.Flex.GCaMP6s.WPRE.SV40, which was a gift from Douglas Kim & GENIE Project (Addgene plasmid #100842).

For retrograde and anterograde tracing, adult mice > P60 were injected. Mice between P24 and P45 were injected for electrophysiological recordings. Mice between P60 and P65 were injected for behavioral experiments.

GCaMP6s-injected mice were implanted with a 400 µm diameter, 0.48 NA photometry fiber with a mirrored tip angled at 45° (MFC_400/430-0.48_4.5mm_MF2.5_MA45, Doric Lenses). The angled tip allowed chronic placement of the fiber in a position that could image the EW without blocking the cerebral aqueduct found immediately dorsal. This was necessary as blocking the cerebral aqueduct leads to high mortality rates. Fiber placement coordinates were +0.5 to 0.6 mm A/P of lambda, ±0.3 to 0.4 mm M/L, and -3.4 to -3.6 mm D/V. Following behavioral experiments, post hoc confirmation of appropriate fiber placement was performed.

### Fixed tissue preparation

Mice were anesthetized with isoflurane and perfused transcardially with 4% paraformaldehyde (PFA) (Electron Microscopy Sciences) in phosphate buffered saline (PBS). Brains and when applicable, spinal columns, were extracted and stored in 4% PFA overnight. Brain slices were made at 80 µm thickness for fluorescent projection mapping and 60 µm for all other analyses. Some brains and all spinal cords were embedded in a gel of 4% low-melting point agarose for slicing (Sigma). All slices were made using a Leica VT1000 S vibratome (Leica Biosystems).

### Immunofluorescence

Slices were stored in PBS and sampled at between 1 in 6 and 1 in 3 for immunofluorescence and expression validation and sampled at 1 in 2 for projection mapping. Immunofluorescence staining protocols varied based on primary antibody. For CART, urocortin, red fluorescent protein, and isolectin B4 staining, slices were permeabilized in 0.2% Triton X-100 for 1-2 hours, blocked with 10% bovine serum albumin (BSA) with 0.05% Triton X-100 for 2 hours, washed, stained with 1:1000 primary antibody (anti-CART Ab, Phoenix Pharmaceuticals, H-003-62; anti-urocortin Ab, Sigma, U4757; anti-RFP Ab, Rockland Immunochemicals, 600-401-379) or 1:1500 biotinylated IB4 (Sigma, L2140) while shaking at 4 °C overnight in solution with 0.2% Triton X-100, washed, stained with 1:500 secondary antibody (Goat Anti-Rabbit 488 or 647, Life Technologies) or 1:500 streptavidin AF488 (Invitrogen S32354), and washed a final time. For choline acetyltransferase (ChAT) staining, the protocol used was from a prior report (Gritton et al., 2016). Briefly, slices were rinsed in Tris-HCL buffer and then blocked and permeabilized in Tris-HCl buffer with 5% donkey serum and 0.2% Triton X-100 for 60 minutes, stained with goat anti-ChAT antibody (AB144P, Millipore) overnight with shaking at 4 °C, rinsed in Tris-HCl buffer with 0.2% Triton X-100, and incubated for two hours in 1:500 secondary antibody (Donkey Anti-Goat 647, Life Technologies). Slices were then mounted on Superfrost Plus microscope slides (Fisher Scientific), dried, and coverslipped with Hoechst 33342 nuclear stain (ThermoFisher Scientific).

### Anatomical imaging

Virally fluorescently expressing brain slices and immunofluorescently labeled brain sections were imaged on an Olympus VS110 imaging system at 10x. Regions of interest were subsequently imaged on a Leica TCS SPE confocal microscope (Leica Microsystems) at 40x. Colocalization counts of immunofluorescence and genetically encoded fluorescent protein signal were performed manually. Quantitative two-dimensional projection mapping was performed using the CCFv3 of the Allen Mouse Brain Atlas (Wang et al., 2020), downloaded in November 2017. Atlas images were manually registered in Adobe Illustrator against raw data images that were obtained by extracting slices imaged on the Olympus VS110 imaging system using the Bioimaging and Optics Platform VSI Reader ActionBar plugin within FIJI (Schindelin et al., 2012). Custom-written MATLAB scripts quantified the number of pixels found with a Sobel edge detector in anatomical regions defined in the atlas.

### Transmission electron microscopy (TEM)

For preparation of TEM tissue, mice were transcardially perfused with ice-cold PBS, followed by ice-cold PBS containing 2% glutaraldehyde and 2% PFA. Following overnight post-fixation at 4 °C in the same fixative, coronal brain slices were made at 100 µm on a Leica VT1000 vibratome. Slices were incubated with 3,3’-Diaminobenzidine (DAB) with metal enhancer (D0426, Sigma). DAB solution (0.25 mg/ml DAB, 0.1 mg/ml CoCl_2_, 0.15 mg/ml H_2_O_2_) was prepared by dissolving DAB and hydrogen peroxide tablets separately in 5 ml of PBS for each. Solutions were mixed immediately before use. Brain slices were incubated in DAB solution for ∼3 min to selectively label APEX-containing cellular structures. After DAB precipitation, slices were washed several times with 0.05 M sodium phosphate buffer (PB), and then processed for TEM with 2 exchanges of fixative that consisted of 2.5% glutaraldehyde, 2% PFA in 0.1 M PB.

Slices were washed 3x with buffer followed by a secondary fixation in 1.5% osmium tetroxide (aqueous). Samples were washed 3x with DI water before beginning an acetone dehydration series. Osmium staining, washes, and acetone dehydration series were carried out in a Pelco Biowave Microwave with Cold Spot and vacuum. EMBed 812 embedding media by EMS was gradually infiltrated with acetone for flat embedding. Selected ROIs were cut out and mounted on a blank stub for sectioning. 90 nm thin sections were collected on copper grids using a Leica Ultracut S ultramicrotome and DiATOME 45° diamond knife. Images were acquired at 100 kV on a 1230 JEOL TEM and Gatan Orius camera with Digital Micrograph software. Magnification for quantified images was 8000x. This work made use of the BioCryo facility of Northwestern University’s NUANCE Center, which has received support from the SHyNE Resource (NSF ECCS-2025633), the IIN, and Northwestern’s MRSEC program (NSF DMR-1720139).

### Fresh tissue preparation

Coronal brain slices were prepared from P40 to P65 mice that had been deeply anesthetized with isoflurane and then perfused transcardially with cold, oxygenated ACSF containing, in mM, 127 NaCl, 2.5 KCl, 25 NaHCO_3_, 1.25 NaH_2_PO_4_, 2 CaCl_2_, 1 MgCl_2_, and 25 glucose with a final osmolarity of around 310 mOsm/L. Extracted brains were sliced in cold ACSF with the support of a small piece of 4% agar. Slices were made at 250 µm thickness for striatum and 300 µm for all others using a Leica VT1000s vibratome and then transferred into a holding chamber with ACSF equilibrated with 95% O_2_/5% CO_2_, where they were incubated at 34°C for 20-30 minutes prior to recording. Slices for recording optogenetic stimulation of EW terminals in the bed nucleus of the stria terminalis were prepared similarly, but with a cold sucrose cutting solution in place of ACSF for the perfusion and slicing steps; slices were then incubated in ACSF at 34°C for 60 minutes prior to recording. Sucrose cutting solution contained, in mM, 194 sucrose, 20 NaCl, 4.4 KCl, 26 NaHCO_3_, 1.2 NaH_2_PO_4_, 2 CaCl_2_, 1 MgCl_2_, and 10 glucose (Kash and Winder, 2006).

### Electrophysiology

The recording chamber was perfused with oxygenated ACSF. Neurons were visualized with QIClick CCD camera (QImaging) under the control of MicroManager (Edelstein et al., 2014), using infrared DODT contrast under a 40x or 60x water-immersion objective (LUMPlan FL, Olympus) with a PE300 CoolLED illumination system (CoolLED Ltd.) providing illumination for fluorescence visualization and optogenetic stimulation, as required. For current clamp experiments, optogenetic stimulation was performed with a 460 nm LED at a power of 10 mW, at 30 Hz with a 10 ms pulse duration (Knobloch et al., 2012). For SPNs, the amplitude of minimal current injection necessary for evoking action potentials was determined manually in intervals of 25 pA. Action potentials were also evoked at 50 pA below and above this value. In the BST, minimal current injection was determined in intervals of 5 pA. For all current clamp optogenetic experiments, current injections were performed with 10 s between current injections. The order of the three amplitudes of injected current was varied randomly between cells. For voltage clamp experiments to measure fast neurotransmitter release, optogenetic stimulation was performed with at a power of 10 mW at 0.5 Hz with a 10 ms pulse duration. Excitatory postsynaptic currents were measured at a holding potential of -70 mV and inhibitory postsynaptic currents were measured at a holding potential of 0 mV. Recordings with a leak current >150 pA for EPSCs and >300 pA for IPSCs were excluded from analysis.

Electrophysiological recordings were obtained using an Axon 700B amplifier (Axon Instruments), sampled at 10-20 kHz, and filtered at 3-5 kHz with ScanImage, an adapted MATLAB-based acquisition package (Pologruto et al., 2003). BNC-2110 data acquisition boards (National Instruments) were used for data acquisition and amplifier and LED pulse control. For whole cell current clamp and cell-attached recordings, the internal solution contained, in mM, 135 K-gluconate, 4 KCl, 10 HEPES, 10 Na_2_- phosphocreatine, 4 MgATP, 0.4 Na_2_GTP, and 0.5 to 1 EGTA (pH 7.2, ∼295 mOsm/L). For some recordings, compounds were added to the internal solution to visualize cell morphology or confirm cell identity and location: Alexa Fluor 488 (10-20 µM, Thermo Fisher Scientific), Neurobiotin (0.1%, Neurobiotin 488 tracer, Vector Laboratories), Neurobiotin-Plus (0.5%, Vector Laboratories). To block Gα_q_ signaling pathways, 10 µM U 73122 (Tocris) was added to the bath solution. To block Gα_s_ signaling pathways, 20 µM PKI (5-24) (Tocris) was added to the internal solution. For rM3Ds validation, currents were recorded from mCherry^+^ EW neurons in cell-attached mode. 10-20 µM clozapine N-oxide (Enzo Life Sciences) dissolved in either ACSF or saline was added to the bath solution. For whole cell voltage clamp recordings, the bath solution contained 5 µM CPP and the internal solution contained, in mM, 120 CsMeSO_3_, 15 CsCl, 10 HEPES, 2 QX-314 Cl, 2 MgATP, 0.3 Na_2_GTP, and 1 EGTA (pH ∼7.2, ∼295 mOsm/L).

Electrophysiological data were analyzed in MATLAB. For measurements of current-evoked action potentials, 0 mV threshold was imposed for identification. Action potentials were averaged across five trials for optogenetic experiments and across three trials for ligand application experiments. For cell-attached mode, action potential current rates were averaged across 60 s. For EPSCs and IPSCs, currents were compared to the 50 ms prior to the light stimulation and the duration of light stimulation combined with the 5 ms after the cessation of the light. The existence of a peak >5 pA away from both these baselines was taken as an EPSC or IPSC. Under this analysis, spontaneous evoked EPSCs and IPSCs were observed throughout the traces. Positive EPSC/IPSC identification was then manually validated based on the latency between the onset of light stimulus and the onset of the PSC. Currents with latencies between 0 and 15 ms were considered ‘evoked’. For quantification of PSC amplitude, the mean of the 15 ms following the onset of the light was compared to the mean of the 15 ms preceding light onset.

### Behavior

Male and female mice were used for all experiments. Experimental animals were injected with AAV1-CBA-DIO-rM3Ds-mCherry-WPRE or AAV1-EF1α-Lox-mCherry-lox(dtA)-lox2. Control animals were injected with volume-matched AAV1-CAG-FLEX-tdTomato-WPRE or AAV9-EF1α-DIO-eYFP-WPRE. In a single cage of littermates, experimental and control animals were counterbalanced. Posthoc validation of viral expression was performed on all experimental animals, and animals lacking signs of expression were excluded. Researchers were blinded to condition during all animal handling and analysis of behavior. Following the surgery, mice were housed for three weeks prior to any behavioral tests or clozapine N-oxide (CNO) administration to allow for sufficient viral expression. Prior to being run in any behavioral tests, the mice were acclimated to handling and the researcher running the behavioral tests. Acclimation consisted of letting the mouse sit in the palm of the researcher’s hand for two to three minutes and took place on two days. Mice that would later be injected with CNO were scruffed briefly (∼10 s) immediately following acclimation, but no injection was given.

Video recordings were made for all behavioral tests except the von Frey test. Cameras were a STC-MC33USB (Sentech) with a YV2.8x2.8LA-2 lens (Fujinion, Fujifilm) with StCamSWare software (Sentech) for the dtA elevated plus maze experiments, a Blackfly S BFS-U3-16S2M-CS (FLIR) with a 3.5-8.0mm F1.4 lens (Yohii, Amazon) with Spinview software (FLIR) for the looming stimulus test, and a Raspberry Pi camera module v2 for all other experiments, acquiring at 25 or 30 frames per second. Recordings were converted from .h264 to .mp4 files with Yamb. For open field locomotion and the elevated plus maze with dtA mice, mouse position was tracked using idTracker (Pérez-Escudero et al., 2014) and custom Matlab scripts were used to analyze behavior. For all other tests, behavior was scored manually by an investigator blinded to the condition of the animal. To minimize circadian influences, behavioral tests were performed at least one hour from zeitgeber, and all animals were run within a 3 hour time window for any given behavioral test. Between individual animals the behavioral setup was cleaned with 70% ethanol, with residual ethanol allowed to dissipate for about five minutes. The one exception was fear conditioning, in which animals were run so that there was consistently 24 hours between fear conditioning and fear retrieval and the behavioral apparatus was cleaned with 1% acetic acid.

For experiments comparing rM3Ds to control injected mice, all mice were given a 3 mg/kg intraperitoneal (i.p.) injection of CNO 40-45 minutes prior to behavioral testing. CNO (Enzo Life Sciences) was diluted in sterile saline to 3 mg/kg for 0.1 mL injected per 20 g. Control mice for rM3Ds experimental groups are given injections during handling, while control mice for dtA experimental groups are not.

#### Fear conditioning

Experiments were performed during the animals’ dark cycle. Mice were acclimated near the behavioral room, outside of the range of olfactory and auditory cues, for >30 minutes prior to injection, and >60 minutes prior to fear testing. Fear conditioning and retrieval, modified from (Senn et al., 2014), was performed in a 19 cm x 19 cm x 27 cm chamber of an Active/Passive Avoidance Shuttle Box (MazeEngineers) with a grid floor that delivers electrical shocks. The room was dark, and the arena was illuminated with red LED bulbs. Prior to each experiment, the floor and behavioral apparatus were cleaned with 1% acetic acid. White paper towels saturated with 1% acetic acid were placed in the bottom of the collecting chamber beneath the box to provide stronger olfactory cues and improve visual contrast. Mice were placed in the box and after two minutes of acclimation were given four cues of a 7.5 kHz tone at 70 dB simultaneous with a dim light cue for 30 s (CS). A 0.5 mA, 1 s shock (US) was delivered immediately following the cessation of each cue. The time between shocks was randomized to 110±50 s. Mice remained in the chamber for 2 additional minutes prior to being gently removed and returned to their home cage. Twenty-four hours later, mice were placed in the box once more. Following five minutes of acclimation, four CS were given, with two minutes between each cue. Fear behaviors were evaluated manually. Freezing bouts were counted when they were ≥2 s in length.

#### Looming stimulus

Mice were acclimated to the behavioral room for at least 30 minutes. Mice were placed in a 30.5 cm x 30.5 cm x 30.5 cm clear plexiglass box with a monitor displaying a gray background mounted above the box. The arena was lit with red light bulbs or infrared LEDs. After five minutes of acclimation to the box, a looming stimulus (Yilmaz and Meister, 2013) on the monitor was triggered when the mouse was underneath the monitor, generally eliciting a freezing response. The looming stimulus was comprised of ten presentations of an expanding black disc; each presentation expanded for 250 ms, held at its maximum size for 250 ms, and then displayed the original gray background for 500 ms. Freezing was evaluated manually during the presentation of the looming stimulus and for the subsequent 30 s (Zelikowsky et al., 2018), using the same parameters as those used for fear conditioning. Experiments were performed during the animals’ dark cycle.

#### Elevated maze

Mice were acclimated near the behavioral room for at least 15 minutes. rM3Ds and dtA mice, with their controls, were placed on a 58 cm x 58 cm elevated plus maze raised 61 cm off the ground. Exploratory behavior was recorded for five minutes. When behavior was analyzed manually, an entrance to an open arm was marked when all four paws were on the open arm. Light intensity was ∼100-150 lux. Experiments were performed during the animals’ dark cycle.

#### Open field locomotion

Mice were acclimated near the behavioral room for at least 15 minutes. rM3Ds mice were placed in the center of a square open field 54 cm on a side with walls 30 cm high. Mice were allowed to explore the box freely for ten minutes. The amount of time spent in the center of the box was analyzed for both the first five minutes and the full ten minutes. The center was defined as either the middle square of the arena when divided into a 3×3 grid, or the four middle squares of the area when divided into a 4×4 grid. The data displayed reflects the amount of time spent in the middle of a 3×3 grid for the first five minutes, but regardless of chosen parameters, no statistically significant differences were seen between control and experimental animals. Light intensity was ∼250 lux. Experiments were performed during the animals’ dark cycle.

#### Light-dark box test

Mice were acclimated to the behavioral testing room for at least 30 minutes. Mice were placed in the center of the bright side of a light-dark box, with the bright side illuminated at ∼350 lux with dimensions of 30.5 cm x 30.5 cm x 30.5 cm and the dark side with dimensions of 15.25 cm x 30.5 cm x 30.5 cm. A hole 4 cm wide by 4.5 cm high connected the two chambers. Mice were allowed to freely explore the box for ten minutes. The amount of time the mouse spent on the bright side was measured visually when all four paws were out of the dark side. Experiments were performed during the animals’ dark cycle.

#### Novelty suppressed feeding

Mice were fasted for 24 hours. Mice were acclimated in the behavioral room for at least 60 minutes. During their dark cycle, mice were placed in an arena 54 cm x 36 cm x 15 cm with bedding, illuminated at ∼800 lux, with a piece of regular chow affixed using a rubber band to a petri dish with filter paper in the center of the arena, following previously developed protocols (Samuels and Hen, 2011). Mice were removed from the arena after ten minutes or after a mouse approached the chow and ate it, whichever came first.

#### von Frey test

Experiments were performed during the animals’ light cycle at ∼210 lux. Following acclimation to the room for at least 30 minutes, animals were acclimated for 30 minutes underneath glass beakers with an 11 cm diameter on a wire mesh. Mice were then given an injection of CNO and placed in their home cage for ten minutes before returning to the glass beakers for 20 minutes. The mechanical sensitivity was tested using von Frey filaments from a Touch Test Sensory Evaluator (Stoelting Physiology Instruments). Sensitivity was determined using the up-down method (François et al., 2017; Peirs et al., 2015), beginning at 1.5 and not going beyond 8. Testing was performed on both left and right paws with five minutes between each test. Paw withdrawal threshold scores for each paw were calculated using the Dixon method (Chaplan et al., 1994) and averaged together for each mouse.

### Photometry recording

Recording of GCaMP6s was performed >6 days after fiber implantation and 3-6 weeks post-viral injection. Excitation was performed with a 470 nm LED or a 405 nm LED at a power of 15-30 µW at the fiber tip (ThorLabs, M470F3, M405FP1) coupled through a fiber optic patch cable (ThorLabs, 200 µm, 0.39 NA) to a 6 port fluorescence Mini-cube (Doric Lenses). Emission was collected through a fiber optic patch cable (Doric Lenses, 400 µm, 0.48 NA) coupled to a rotary joint commutator (Doric Lenses). A fiber optic patch cable (ThorLabs, 600 µm, 0.48 NA) transmitted emission light to the detector (Wu et al., 2021b). Emission was detected with a Newport visible femtowatt photoreceiver (Doric Lenses) and collected at 250 Hz through a BNC-2110 data acquisition board using electrophysiology acquisition scripts. Acquisition of fluorescence triggered a simultaneous 25 fps video recording with a Raspberry Pi camera module v2. Behavioral recordings were taken from a square open field box, the mouse home cage, an isoflurane induction chamber, or from one side of an Active/Passive Avoidance Shuttle Box (MazeEngineers).

Spontaneous slipping was taken from 4 to 6 slipping events for each mouse at 470 nm excitation. Tail restraint was performed for 8-10 s, 5 to 7 times for each mouse at 470 nm excitation and once at 405 nm excitation. Anesthetic induction and recovery were performed 3 to 4 times for each mouse at 470 nm excitation and once at 405 nm excitation. Tail suspension was performed for ∼ 6 s, 6 to 9 times for each mouse at 470 nm excitation and 3 times at 405 nm excitation. Looming stimuli were presented by a dark paper bird shape repeatedly moved downwards towards the mouse for ∼10 s with an interstimulus interval of ∼1 s. Each looming stimulus trial was given 6 to 7 times to each mouse at 470 nm excitation and once at 405 nm excitation. Stressful white noise stimulus was given at ∼90 dB for 10 s (Kim et al., 2019). White noise was presented 6 to 8 times at 470 nm excitation and once at 405 nm excitation.

For shock and fear responses, an acoustic cue (15 s, 7.5 kHz tone at 70 dB) was given 3 to 6 times. This acoustic cue was then paired with a 1 s electric shock (0.6 mA) immediately following the cessation of the tone. 8 cue-shock pairs were given, with 7 presentations at 470 nm excitation and 1 at 405 nm excitation. The acoustic cue was then given in the absence of the shock an additional three times at 470 nm excitation. Analysis of fear photometry was averaged from the final three cue-shock pair trials and the three cue-only trials following fear conditioning. For all others, fluorescence responses were averaged across trials for each mouse. Fluorescence traces were generally aligned to the time-locking stimulus and baselined using the time period 20 s to 10 s prior to the stimulus. For motor dysfunction following recovery from anesthesia, traces were baselined using the time period 5 s to 0 s prior to the stimulus. Peak z-scores were calculated from the average of 250 values surrounding the maximum found during the baseline and during the stimulus. For baseline behavior recordings, in the absence of extraneous stimuli, 8 minutes of behavior were acquired. Z-scores were calculated across the recordings and thresholded at a z-score of 3 for at least 100 ms. Analysis was performed in MATLAB.

### Data display

All behavioral schematics depicted in Figures 5, 6, 8, S6, and S8 were created with BioRender.com. Whole brain RNAseq data (Zeisel et al., 2018) was downloaded from wholebrain.org and analyzed using custom MATLAB scripts.

### Statistical analyses

Statistical analyses were performed in Prism 8.4 (GraphPad Software). All statistical tests were two-sided. Statistical details of experiments can be found in the figure legends. Data are reported as mean ± SEM.

---

**Figure S1:**
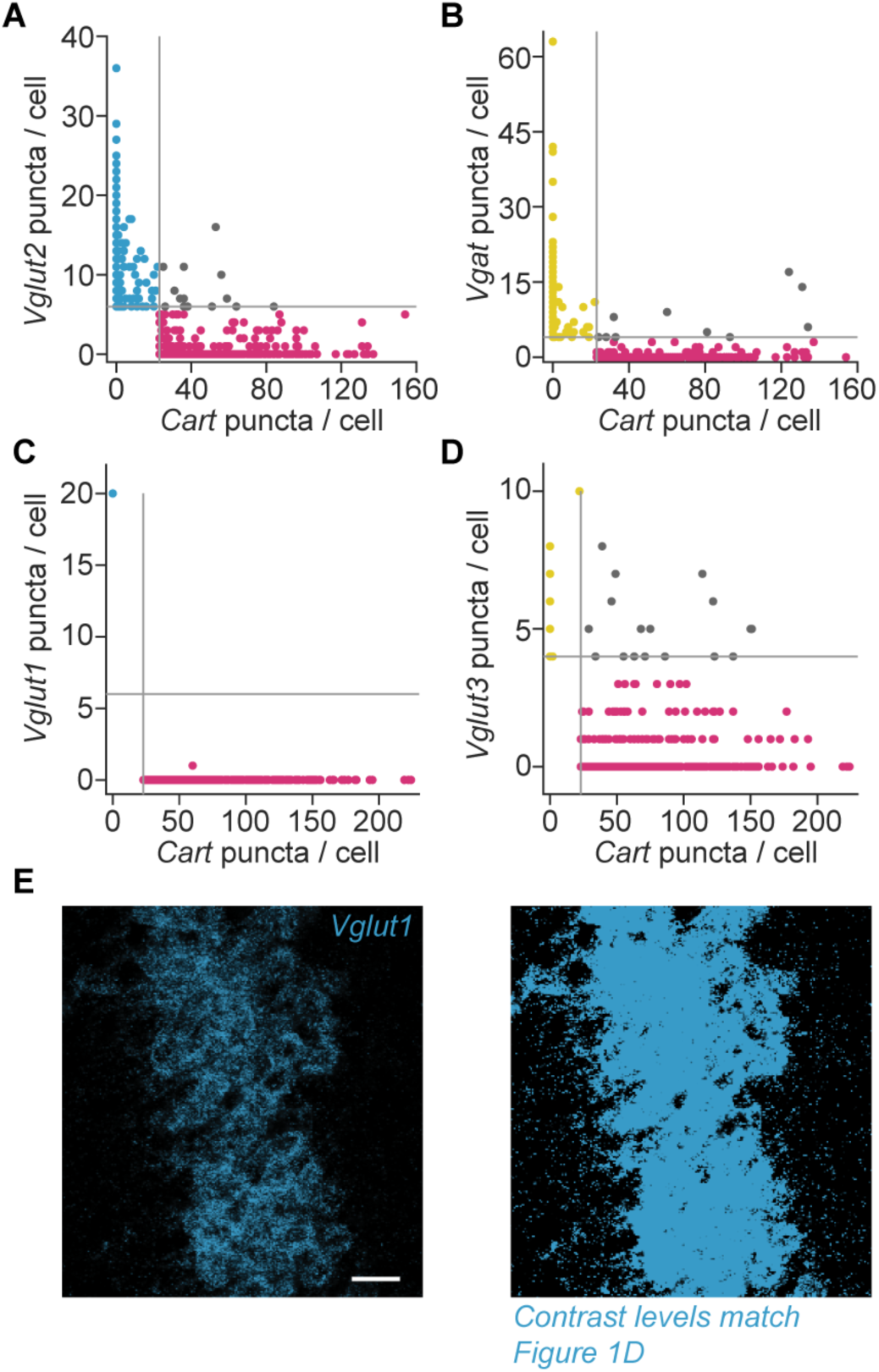
Quantification of vesicular transporter transcripts. (A) Quantification of puncta per cell in the EW and peri-EW regions for *Vglut2*/*Slc17a6* (blue, 744 neurons), and *Cart*/*Cartpt* (red, 297 neurons). Gray lines denote thresholds above which neurons were considered positive for the transcript, and gray dots are neurons positive for both transcripts (15 neurons). Data are pooled from 3 mice. (B) As in (A) but for *Vgat*/*Slc32a1* (yellow, 249 neurons). 10 neurons are positive for both. (C) Similar to (A), but for *Vglut1*/*Slc17a7* (blue, 1 neuron) against *Cart* neurons (red, 404 neurons). 0 neurons are positive for both. (D) As in (C), but for *Vglut3*/*Slc17a8* (yellow, 28 neurons). 17 neurons are positive for both. (E) Robust *Vglut1* transcript signal is seen in the hippocampus (left; scale, 20 µm). The same image is shown on the right, with contrast levels set identical to those used in the main text for *Vglut1*. Absence of *Vglut1*/*Slc17a7*-positive neurons in and near the EW is not due to insufficient probe labeling.

**Figure S2:**
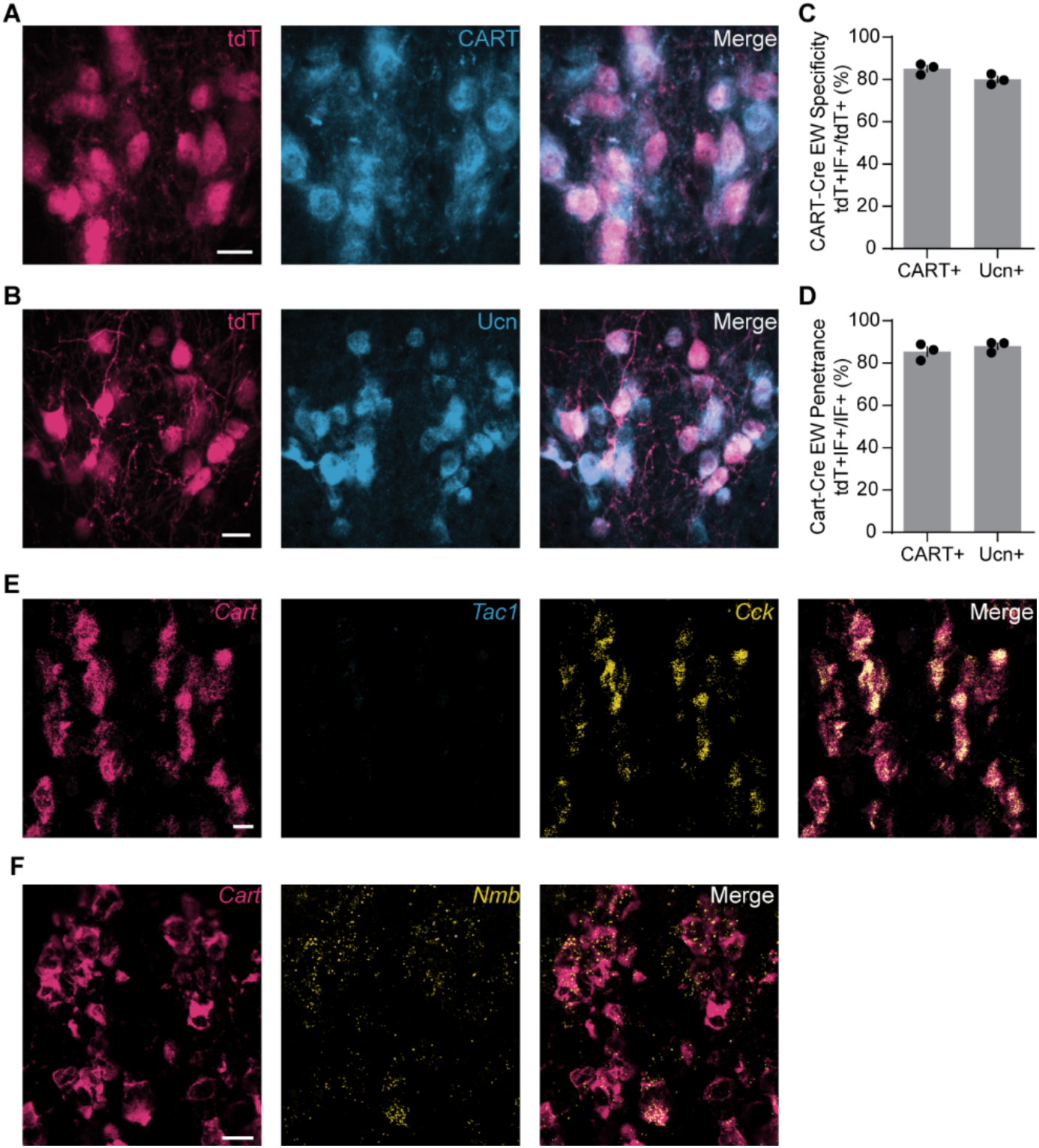
Genetic targeting and neuropeptides of the CART^+^ EW. (A) tdT^+^ neurons of the EW (left, red) compared to immunofluorescence against CART peptide (middle, blue) show high levels of colocalization (right, merge, magenta). Scale, 20 µm. (B) As in (A) but with immunofluorescence against urocortin (blue). (C) Quantification of tdT^+^ EW cells positive for CART (85.1%±1.5%, n=3 mice, 784 cells) and urocortin (Ucn, 80.0%±1.4%, n=3 mice, 631 cells). (D) Quantification of CART^+^ (85.5%±2.3%, n=3 mice, 677 cells) and Ucn^+^ (88.0%±1.5%, n=3 mice, 574 cells) that are tdT^+^. (E) FISH against *Cart* (red), *Tac1*/*Sub. P* (blue) and *Cck* (yellow). As seen in the merge, there is robust colocalization between *Cck* and *Cart*. Scale, 20 µm. (F) Similar to (E), but *Cart* (red) and *Nmb* (yellow) showing colocalization in some EW neurons (merge). Error bars represent SEM.

**Figure S3:**
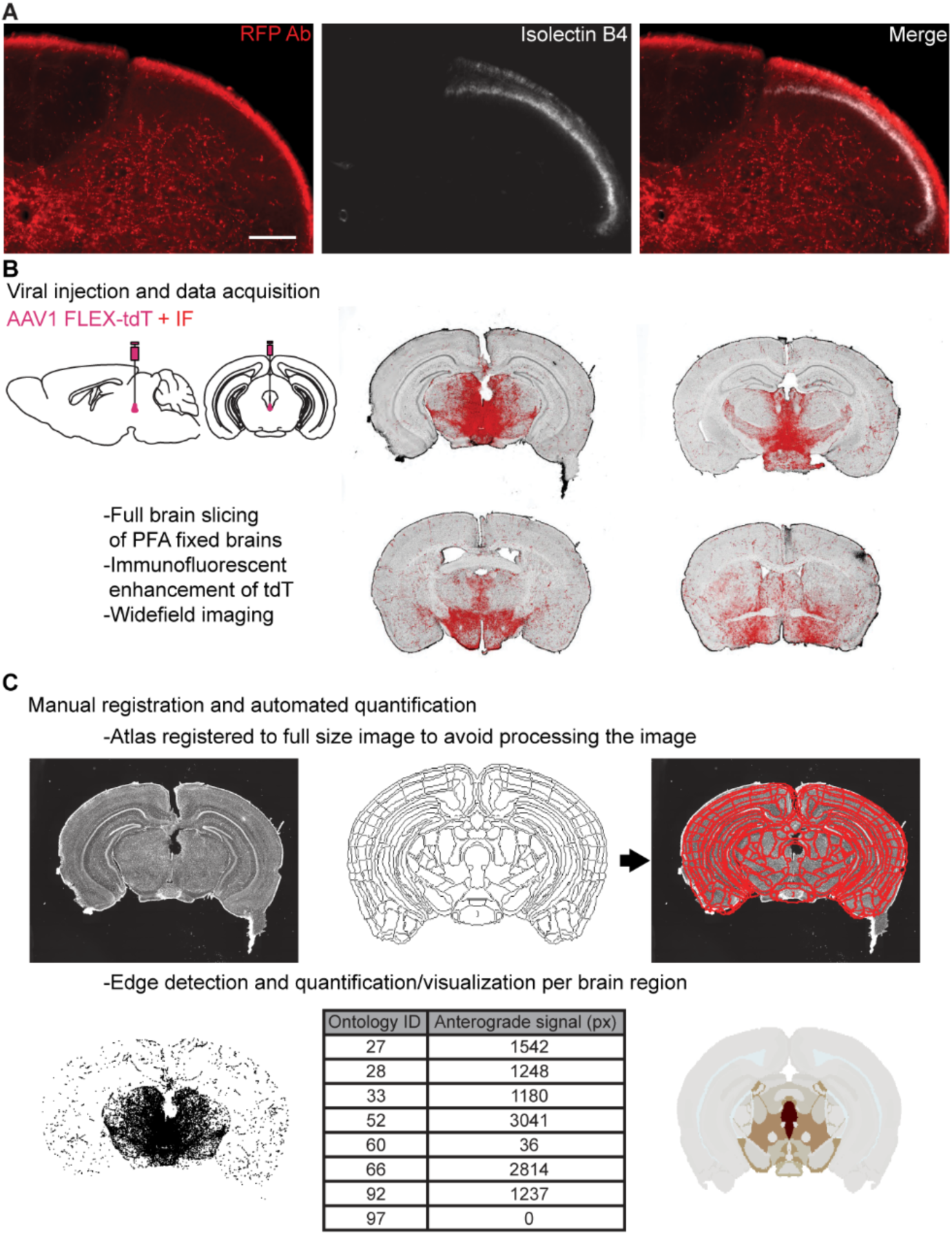
Spinal cord lamina labeling and whole brain mapping of CART^+^ EW projections. (A) CART^+^ EW projections (red) in the spinal cord do not appear to extend to lamina II, marked with isolectin B4 (white), but are found in the pain- and mechanically-sensitive dorsal horn and lamina X. Scale bar, 100 µm. (B) Visualization of the whole brain mapping pipeline. Injection of AAV FLEX-tdTomato in the CART^+^ EW was followed by fixed slicing, immunofluorescent enhancement of tdTomato, and widefield fluorescent imaging to produce whole brain sections (set of 4 is shown here as an example). (C) Brain sections (top left) were manually registered to coronal slices from the Allen Brain Atlas (top, middle; registration overlay at top, right). Edge detection (bottom, left) was performed to find neuronal processes throughout the brain section. Processes were quantified for each brain region (bottom, middle) and visualized as a heatmap.

**Figure S4:**
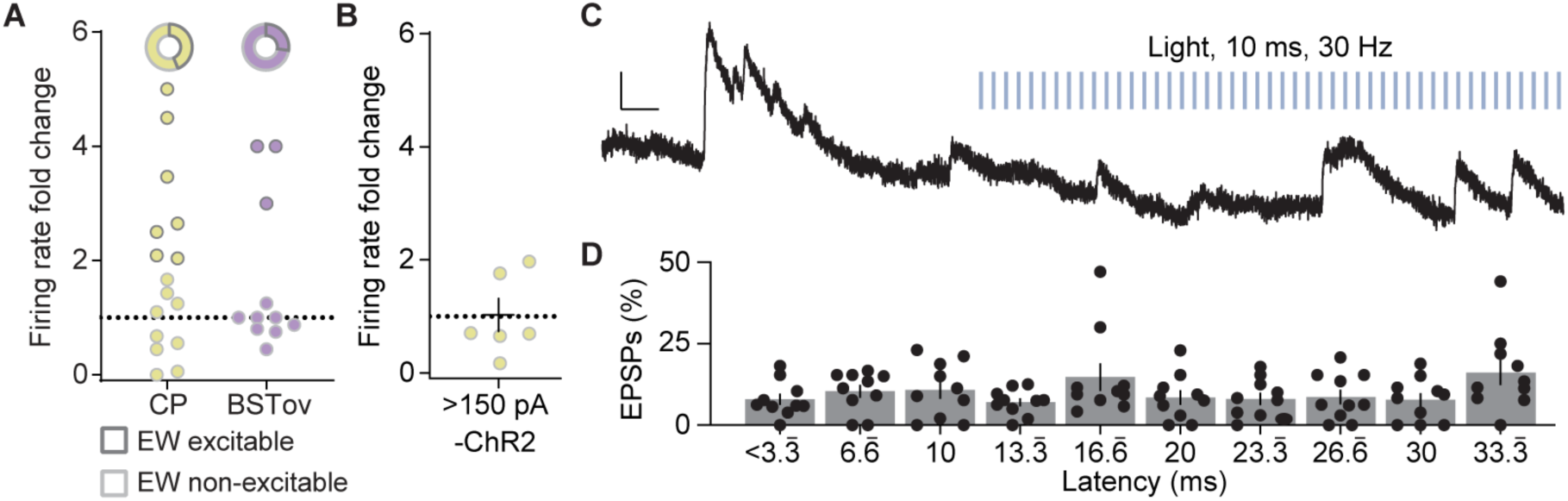
Functional peptidergic projections of the CART^+^ EW. (A) Firing rate fold change of neurons recorded from CP (yellow) and BSTov (purple). Below, neurons in each region increase firing rate >2x following optical stimulation (dark gray, 7 of 16 SPNs in CP; 3 of 11 BST neurons). (B) CP SPNs with rheobase >150 pA did not increase to light stimulation in the absence of CART^+^ EW ChR2. p = 0.9367, paired t-test, t_(5)_ = 0.0834, n = 6 neurons. (C) Optogenetic stimulation did not elicit EPSPs in neurons activated by ChR2 CART^+^ EW fibers. Blue lines denote light pulses. Scale bars, 0.5 mV, 100 ms. (D) Quantification of EPSP latencies (ms) binned by 3.3 ms across the 33.33 ms interstimulus interval. No bin had a higher proportion of EPSPs than any other (p = 0.2930, repeated measures one-way ANOVA F_(2.555, 22.99)_ = 1.312, n = 10 neurons). Error bars represent SEM.

**Figure S5:**
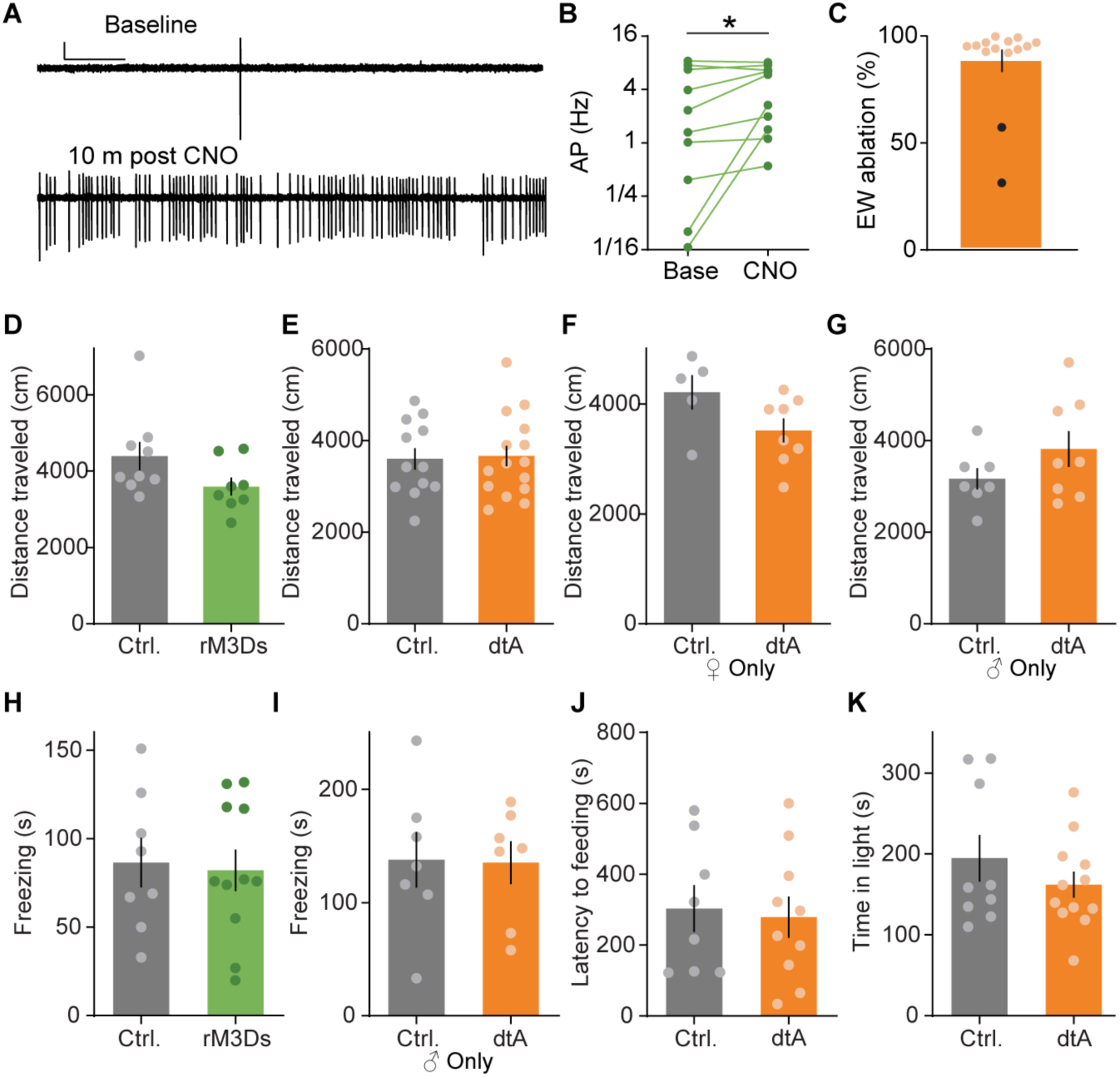
CART^+^ EW neurons selectively promote fear of pain. (A) Cell-attached recording of an rM3Ds-expressing EW neuron shows robust activation after CNO application. Scale, 50 pA, 10 s. (B) Summary data of rM3Ds activation of CART^+^ EW (n = 10 neurons, *p<0.05, paired t-test, t_(9)_ = 2.333, p = 0.0445). (C) Example of quantification of EW ablation following dtA expression. Injection of dtA routinely killed >80% of neurons (orange). Mice with many EW cells remaining (black, n = 2)) were excluded from behavioral analysis. (D) The distance traveled in an open field does not differ between control and rM3Ds-expressing mice (t_(15)_ = 1.768, p = 0.0974, n = 9 Ctrl. mice, 8 rM3Ds mice). p > 0.05, unpaired t-test for all behavioral comparisons. (E-G) As in (D), but comparing control and dtA expressing mice of (E) both sexes (t_(26)_ = 0.1995, p = 0.8434, n = 12 Ctrl. mice, 16 dtA mice), (F) females only (t_(11)_ = 1.885, p = 0.0860, n = 5 Ctrl. mice, 8 dtA mice), and (G) males only (t_(13)_ = 1.372, p = 0.1933, N = 7 Ctrl. mice, 8 dtA mice). (H) Contextual fear of control versus rM3Ds-expressing mice (t_(17)_ = 0.2422, p = 0.8115, n = 8 Ctrl. mice, 11 rM3Ds mice). (I) As in (H) but control versus dtA-expressing male mice (t_(12)_ = 0.0782, p = 0.9390, n = 7 Ctrl. mice, 7 dtA mice). (J) Latency to feeding is unchanged between dtA and control mice in a novelty-suppressed feeding test (t_(16)_ = 0.2732, p = 0.7882, n = 8 Ctrl mice, 10 dtA mice). (K) Time spent on the light side of a light-dark box is unchanged between dtA and control mice (t_(19)_ = 1.058, p = 0.3034, n = 9 Ctrl. mice, 12 dtA mice). Error bars represent SEM.

**Figure S6:**
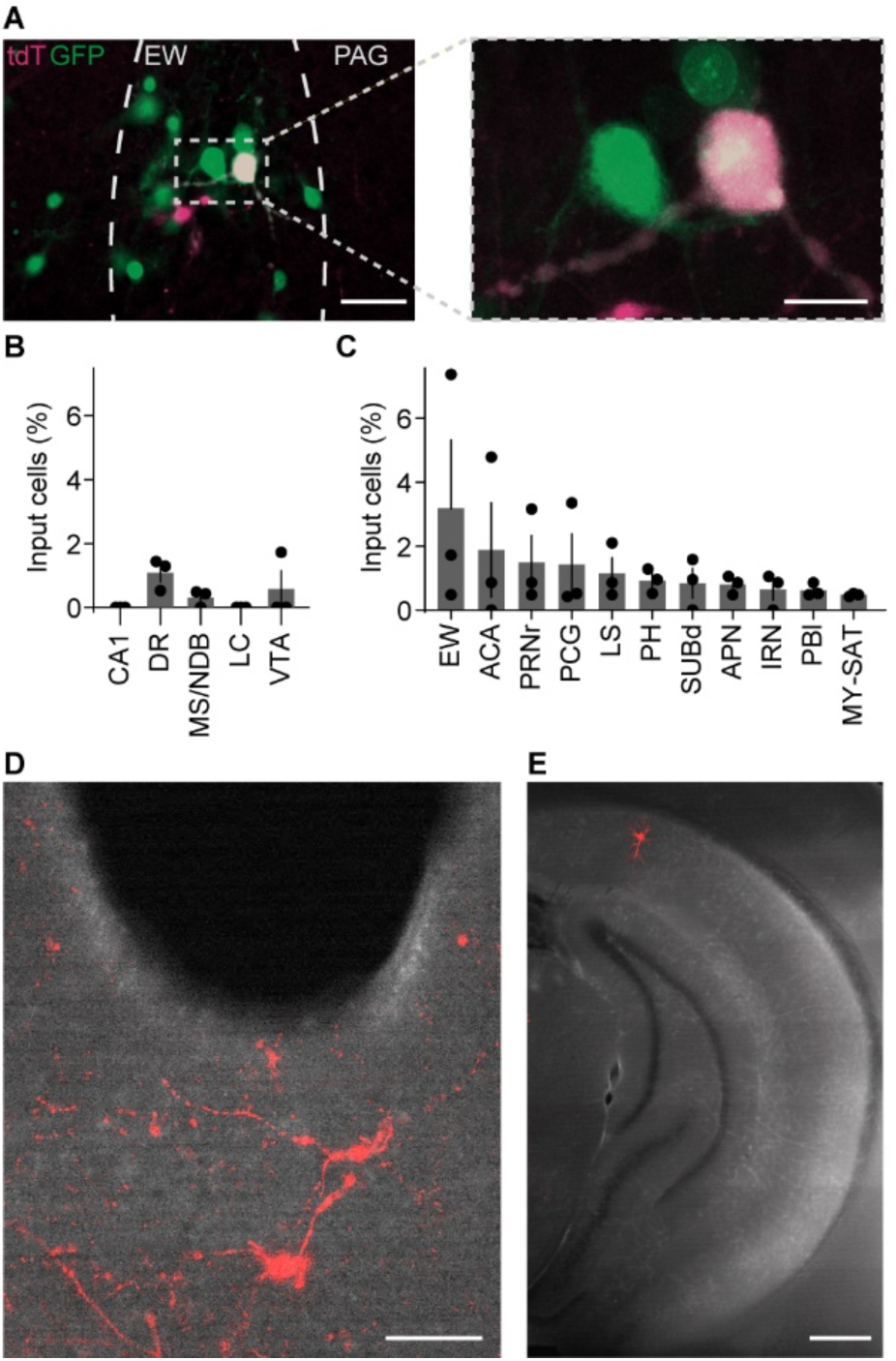
Monosynaptic retrograde tracing reveals a novel collection of inputs to the CART^+^ EW. (A) Fluorescence image (left: scale, 60 µm) of EW injected with CVS-N2c-ΔG-EnvA-tdT rabies virus (red) and CVS-N2c-ΔG- EnvA rabies virus helper viruses (green). Dashed gray box denotes inset shown in confocal image on right (Scale, 20 µm). Individual starter cells can be seen. (B) Monosynaptic retrograde labeling in brain regions previously proposed to project to the CART^+^ EW, as a percentage of total retrogradely labeled neurons (n = 3 mice). These areas show little to no evidence of monosynaptic retrograde labeling. (C) As in (B), but brain regions with variable or sparse monosynaptic retrograde expression, as a percentage of total retrogradely labeled neurons (n=3 mice). (D) Widefield image of immunoenhanced tdT^+^ cells (red) in the dorsal raphe. Scale, 100 µm. (E) Similar to (E), but for the hippocampus. No retrogradely labeled neurons were observed in the hippocampus except for a sparse population in pyramidal neurons of the dorsal subiculum, i.e., SUBd in (C). Scale, 300 µm. Error bars represent SEM.

**Figure S7:**
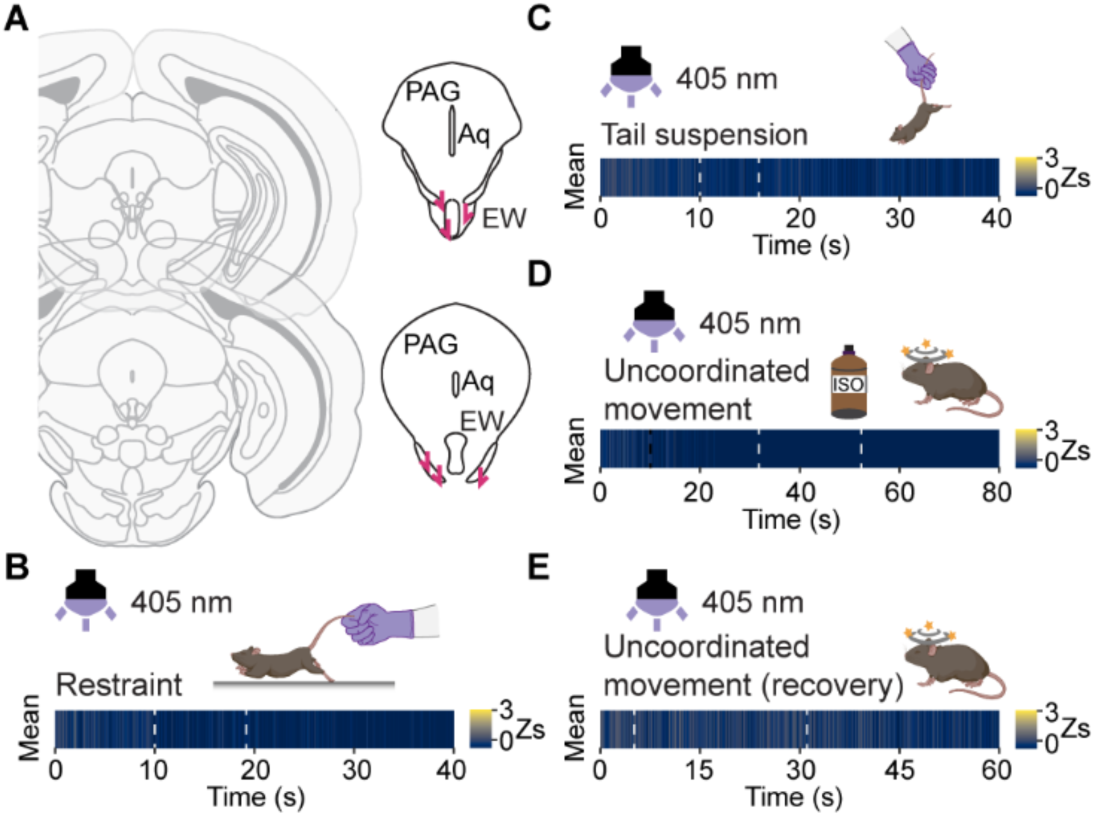
CART^+^ EW GCaMP6s fluorescence signals are not motion artifacts. (A) Schematic of fiber placements. (B) Absence of time-locked fluorescence signals, as in the bottom panel of Figure 7C, in response to tail restraint (n = 6 mice, 1 trial) during excitation at the isosbestic point (purple light, 405 nm). Heatmap is identical to that used in Figures 7E, 7H, and 7I. (C-E) As in (B), with isosbestic excitation, in response to (C) tail suspension (n = 6 mice, 3 trials), (D) uncoordinated movement following anesthetic exposure (n = 6 mice, 1 trial), and (E) uncoordinated movement during recovery from anesthesia (n = 6 mice, 1 trial). Heatmap is identical to that used in (B) and Figures 7E, 7H, and 7I.

**Figure S8:**
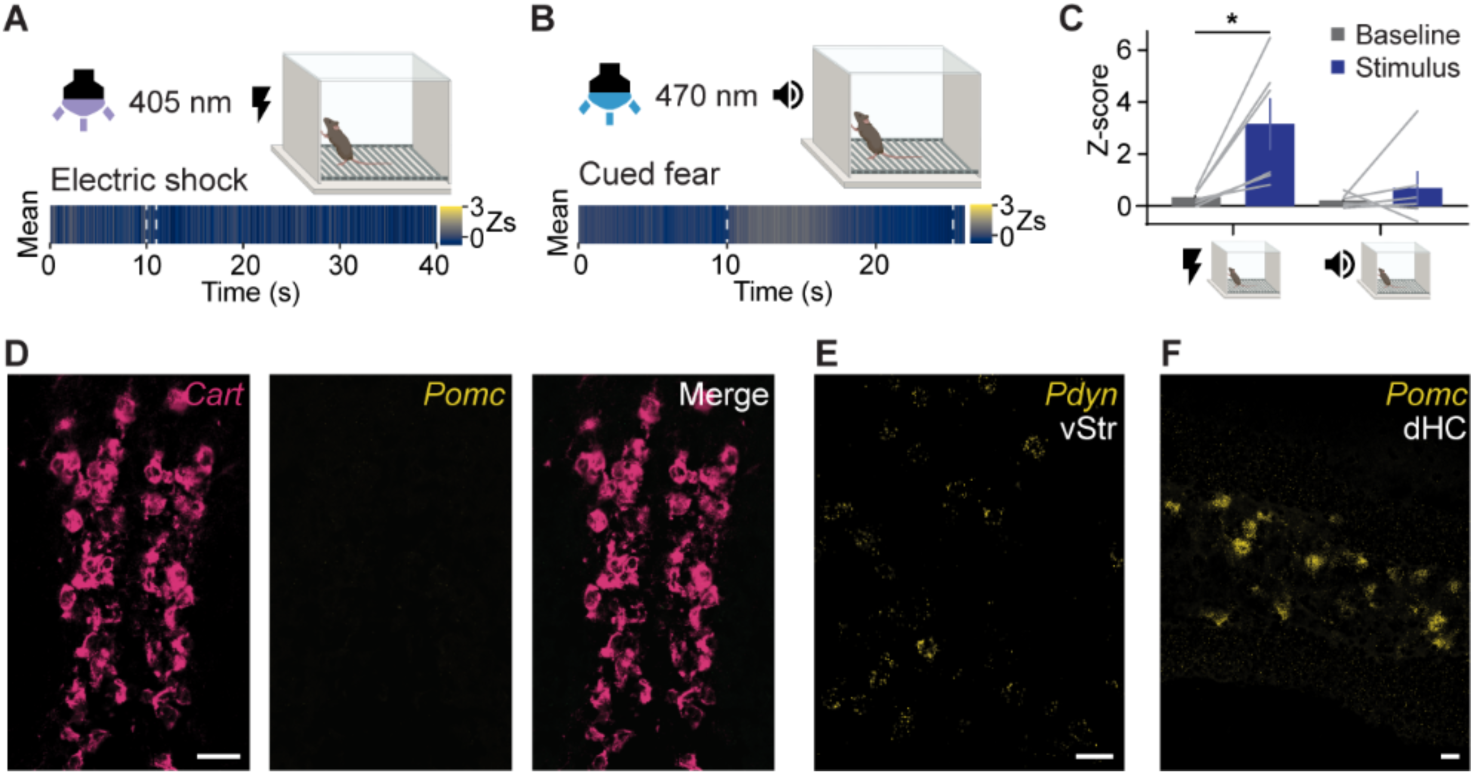
CART^+^ EW neurons respond to pain and lack opioids. (A) Fiber photometry of CART^+^ EW GCaMP6s excited at the isosbestic point shows no fluorescence signal in response to electric shock. Mean from 6 mice, 1 trial per mouse. Gray dashed line marks shock. (B) Exposure to an auditory conditioned fear stimulus did not elicit a consistent fluorescence change from GCaMP6s in the CART^+^ EW. Gray dashed line marks the conditioned stimulus tone. Mean from 6 mice, 6 trials per mouse. (C) Peak z-score of stimulus-driven fluorescence changes versus baseline for shock (left) (*p < 0.05, paired t-test, t_(5)_ = 3.229, p = 0.0232) and fear cue (right) (p = 0.56, Wilcoxon matched-pairs signed rank test). For both stimuli n = 6 mice. (D) FISH against *Cart* (red; scale bar, 20 µm) and *Pomc* (yellow) shows no colocalization in CART^+^ EW neurons (merge). (E) Absence of *Pdyn* neurons in and near the EW is not due to poor probe labeling. FISH against *Pdyn* (yellow) shows clear signal in the ventral striatum. Scale bar, 20 µm. (F) As in (E), but against *Pomc* (yellow) in the dorsal hippocampus. Error bars represent SEM.

## References

Ailion, M., Hannemann, M., Dalton, S., Pappas, A., Watanabe, S., Hegermann, J., Liu, Q., Han, H.-F., Gu, M., Goulding, M.Q., et al. (2014). Two Rab2 Interactors Regulate Dense-Core Vesicle Maturation. Neuron 82, 167–180.

Atasoy, D., Aponte, Y., Su, H.H., and Sternson, S.M. (2008). A FLEX switch targets Channelrhodopsin-2 to multiple cell types for imaging and long-range circuit mapping. J. Neurosci. 28, 7025–7030.

Beinfeld, M., Chen, Q., Gao, F., Liddle, R.A., Miller, L.J., and Rehfeld, J. (2019). Cholecystokinin receptors (version 2019.4) in the IUPHAR/BPS Guide to Pharmacology Database. IUPHAR/BPS Guid. to Pharmacol. CITE 2019.

Bittencourt, J.C., Vaughan, J., Arias, C., Rissman, R.A., Vale, W.W., and Sawchenko, P.E. (1999). Urocortin Expression in Rat Brain: Evidence Against a Pervasive Relationship of Urocortin-Containing Projections With Targets Bearing Type 2 CRF Receptors. J. Comp. Neurol. 415, 285–312.

Bowers, M.E., Choi, D.C., and Ressler, K.J. (2012). Neuropeptide regulation of fear and anxiety: Implications of cholecystokinin, endogenous opioids, and neuropeptide Y. Physiol. Behav. 107, 699–710.

Burnell, J., Ng, L., and Guillozet-Bongaarts, A. (2008). Edinger-Westphal Nucleus. Nat. Preced.

Calhoon, G.G., and Tye, K.M. (2015). Resolving the neural circuits of anxiety. Nat. Neurosci. 18, 1394–1404.

Causeret, F., Moreau, M.X., Pierani, A., and Blanquie, O. (2021). The multiple facets of Cajal-Retzius neurons. Development 148.

Chang, C.-W., Hsiao, Y.-T., and Jackson, M.B. (2021). Synaptophysin Regulates Fusion Pores and Exocytosis Mode in Chromaffin Cells. J. Neurosci. 41, 3563 LP – 3578.

Chaplan, S.R.R., Bach, F.W.W., Pogrel, J.W.W., Chung, J.M.M., and Yaksh, T.L.L. (1994). Quantitative assessment of tactile allodynia in the rat paw. J. Neurosci. Methods 53, 55–63.

Chen, L., Cai, P., Wang, R.-F., Lu, Y.-P., Chen, H.-Y., Guo, Y.-R., Huang, S.-N., Hu, L.-H., Chen, J., Zheng, Z.-H., et al. (2020). Glutamatergic lateral hypothalamus promotes defensive behaviors. Neuropharmacology 178, 108239.

Chen, T.-W., Wardill, T.J., Sun, Y., Pulver, S.R., Renninger, S.L., Baohan, A., Schreiter, E.R., Kerr, R.A., Orger, M.B., Jayaraman, V., et al. (2013). Ultrasensitive fluorescent proteins for imaging neuronal activity. Nature 499, 295–300.

Chou, X., Wang, X., Zhang, Z., Shen, L., Zingg, B., Huang, J., Zhong, W., Mesik, L., Zhang, L.I., and Tao, H.W. (2018). Inhibitory gain modulation of defense behaviors by zona incerta. Nat. Commun. 9, 1151.

Daigle, T.L., Madisen, L., Hage, T.A., Valley, M.T., Knoblich, U., Larsen, R.S.R., Takeno, M.M., Huang, L., Gu, H., Larsen, R.S.R., et al. (2018). A Suite of Transgenic Driver and Reporter Mouse Lines with Enhanced Brain-Cell-Type Targeting and Functionality. Cell 174, 465–480.e22.

Daniel, S.E., and Rainnie, D.G. (2016). Stress Modulation of Opposing Circuits in the Bed Nucleus of the Stria Terminalis. Neuropsychopharmacology 41, 103–125.

Dautzenberg, F.M., Grigoriadis, D.E., Hauger, R.L., Risbrough, V.B., Steckler, T., Vale, W.W., and Valentino, R.J. (2019). Corticotropin-releasing factor receptors (version 2019.4) in the IUPHAR/BPS Guide to Pharmacology Database. IUPHAR/BPS Guid. to Pharmacol. CITE 2019.

Delevich, K., Tucciarone, J., Huang, Z.J., and Li, B. (2015). The mediodorsal thalamus drives feedforward inhibition in the anterior cingulate cortex via parvalbumin interneurons. J. Neurosci. 35, 5743–5753.

Dicken, M.S., Tooker, R.E., and Hentges, S.T. (2012). Regulation of GABA and Glutamate Release from Proopiomelanocortin Neuron Terminals in Intact Hypothalamic Networks. J. Neurosci. 32, 4042 LP – 4048.

Ding, K., Han, Y., Seid, T.W., Buser, C., Karigo, T., Zhang, S., Dickman, D.K., and Anderson, D.J. (2019). Imaging neuropeptide release at synapses with a genetically engineered reporter. Elife 8, e46421.

Dumrongprechachan, V., Soto, G., MacDonald, M.L., and Kozorovitskiy, Y. (2021). Cell type and subcellular compartment specific APEX2 proximity labeling proteomics in the mouse brain. BioRxiv 2021.04.08.439091.

Edelstein, A.D., Tsuchida, M.A., Amodaj, N., Pinkard, H., Vale, R.D., and Stuurman, N. (2014). Advanced methods of microscope control using μManager software. J. Biol. Methods 1, e10.

Fahrenkrug, J., Goetzl, E.J., Gozes, I., Harmar, A., Laburthe, M., May, V., Pisegna, J.R., Said, S.I., Vaudry, D., Vaudry, H., et al. (2019). VIP and PACAP receptors (version 2019.4) in the IUPHAR/BPS Guide to Pharmacology Database. IUPHAR/BPS Guid. to Pharmacol. CITE 2019.

François, A., Low, S.A., Sypek, E.I., Christensen, A.J., Sotoudeh, C., Beier, K.T., Ramakrishnan, C., Ritola, K.D., Sharif-Naeini, R., Deisseroth, K., et al. (2017). A Brainstem-Spinal Cord Inhibitory Circuit for Mechanical Pain Modulation by GABA and Enkephalins. Neuron 93, 822–839.e6.

Gaszner, B., Csernus, V., and Kozicz, T. (2004). Urocortinergic neurons respond in a differentiated manner to various acute stressors in the Edinger-Westphal nucleus in the rat. J. Comp. Neurol. 480, 170–179.

George, D.T., Ameli, R., and Koob, G.F. (2019). Periaqueductal Gray Sheds Light on Dark Areas of Psychopathology. Trends Neurosci. 42, 349–360.

Giardino, W.J., Rodriguez, E.D., Smith, M.L., Ford, M.M., Galili, D., Mitchell, S.H., Chen, A., and Ryabinin, A.E. (2017). Control of chronic excessive alcohol drinking by genetic manipulation of the Edinger–Westphal nucleus urocortin-1 neuropeptide system. Transl. Psychiatry 7, e1021– e1021.

Granger, A.J., Wang, W., Robertson, K., El-Rifai, M., Zanello, A.F., Bistrong, K., Saunders, A., Chow, B.W., Nuñez, V., Turrero García, M., et al. (2020). Cortical ChAT+ neurons co-transmit acetylcholine and GABA in a target- and brain-region-specific manner. Elife 9, e57749.

Grinevich, V., and Ludwig, M. (2021). The multiple faces of the oxytocin and vasopressin systems in the brain. J. Neuroendocrinol., e13004.

Gritton, H.J., Howe, W.M., Mallory, C.S., Hetrick, V.L., Berke, J.D., and Sarter, M. (2016). Cortical cholinergic signaling controls the detection of cues. Proc. Natl. Acad. Sci. 113, E1089–E1097.

Gross, C.T., and Canteras, N.S. (2012). The many paths to fear. Nat. Rev. Neurosci. 13, 651–658.

Hammack, S.E., and May, V. (2015). Pituitary Adenylate Cyclase Activating Polypeptide in Stress-Related Disorders: Data Convergence from Animal and Human Studies. Biol. Psychiatry 78, 167–177.

Han, S., Soleiman, M.T., Soden, M.E., Zweifel, L.S., and Palmiter, R.D. (2015). Elucidating an Affective Pain Circuit that Creates a Threat Memory. Cell 162, 363–374.

Hasan, M.T., Althammer, F., Silva da Gouveia, M., Goyon, S., Eliava, M., Lefevre, A., Kerspern, D., Schimmer, J., Raftogianni, A., Wahis, J., et al. (2019). A Fear Memory Engram and Its Plasticity in the Hypothalamic Oxytocin System. Neuron 103, 133–146.e8.

Hnasko, T.S., Chuhma, N., Zhang, H., Goh, G.Y., Sulzer, D., Palmiter, R.D., Rayport, S., and Edwards, R.H. (2010). Vesicular Glutamate Transport Promotes Dopamine Storage and Glutamate Corelease In Vivo. Neuron 65, 643–656.

Hökfelt, T., Barde, S., Xu, Z.-Q.D., Kuteeva, E., Rüegg, J., Le Maitre, E., Risling, M., Kehr, J., Ihnatko, R., and Theodorsson, E. (2018). Neuropeptide and small transmitter coexistence: fundamental studies and relevance to mental illness. Front. Neural Circuits 12, 106.

Huang, K.W., Ochandarena, N.E., Philson, A.C., Hyun, M., Birnbaum, J.E., Cicconet, M., and Sabatini, B.L. (2019). Molecular and anatomical organization of the dorsal raphe nucleus. Elife 8, e46464.

Innis, R.B., and Aghajanian, G.K. (1986). Cholecystokinin-containing and nociceptive neurons in rat edinger-westphal nucleus. Brain Res. 363, 230–238.

Iwamoto, S., Tamura, M., Sasaki, A., and Nawano, M. (2021). Dynamics of neuronal oscillations underlying nociceptive response in the mouse primary somatosensory cortex. Sci. Rep. 11, 1667.

Kash, T.L., and Winder, D.G. (2006). Neuropeptide Y and corticotropin-releasing factor bi-directionally modulate inhibitory synaptic transmission in the bed nucleus of the stria terminalis. Neuropharmacology 51, 1013–1022.

Keay, K.A., and Bandler, R. (2015). Periaqueductal Gray. Rat Nerv. Syst. Fourth Ed. 207–221.

Kim, J., Zhang, X., Muralidhar, S., LeBlanc, S.A., and Tonegawa, S. (2017). Basolateral to Central Amygdala Neural Circuits for Appetitive Behaviors. Neuron 93, 1464–1479.e5.

Kim, J.S., Han, S.Y., and Iremonger, K.J. (2019). Stress experience and hormone feedback tune distinct components of hypothalamic CRH neuron activity. Nat. Commun. 10, 5696.

Kim, T., Gondré-Lewis, M.C., Arnaoutova, I., and Loh, Y.P. (2006). Dense-Core Secretory Granule Biogenesis. Physiology 21, 124–133.

Knobloch, H.S., Charlet, A., Hoffmann, L.C., Eliava, M., Khrulev, S., Cetin, A.H., Osten, P., Schwarz, M.K., Seeburg, P.H., Stoop, R., et al. (2012). Evoked Axonal Oxytocin Release in the Central Amygdala Attenuates Fear Response. Neuron 73, 553–566.

Kozicz, T. (2003). Neurons colocalizing urocortin and cocaine and amphetamine-regulated transcript immunoreactivities are induced by acute lipopolysaccharide stress in the Edinger-Westphal nucleus in the rat. Neuroscience 116, 315–320.

Kozicz, T., Li, M., and Arimura, A. (2001). The activation of urocortin immunoreactive neurons in the Edinger-Westphal nucleus following acute pain stress in rats. Stress 4, 85–90.

Kozicz, T., Bittencourt, J.C., May, P.J., Reiner, A., Gamlin, P.D.R., Palkovits, M., Horn, A.K.E., Toledo, C.A.B., and Ryabinin, A.E. (2011). The Edinger-Westphal nucleus: A historical, structural, and functional perspective on a dichotomous terminology. J. Comp. Neurol. 519, 1413–1434.

Krashes, M.J., Shah, B.P., Madara, J.C., Olson, D.P., Strochlic, D.E., Garfield, A.S., Vong, L., Pei, H., Watabe-Uchida, M., Uchida, N., et al. (2014). An excitatory paraventricular nucleus to AgRP neuron circuit that drives hunger. Nature 507, 238–242.

Lackey, E.P., Heck, D.H., and Sillitoe, R. V (2018). Recent advances in understanding the mechanisms of cerebellar granule cell development and function and their contribution to behavior. F1000Research 7.

Lefler, Y., Campagner, D., and Branco, T. (2020). The role of the periaqueductal gray in escape behavior. Curr. Opin. Neurobiol. 60, 115–121.

Lein, E.S., Hawrylycz, M.J., Ao, N., Ayres, M., Bensinger, A., Bernard, A., Boe, A.F., Boguski, M.S., Brockway, K.S., Byrnes, E.J., et al. (2007). Genome-wide atlas of gene expression in the adult mouse brain. Nature 445, 168–176.

Li, X., Chen, W., Pan, K., Li, H., Pang, P., Guo, Y., Shu, S., Cai, Y., Pei, L., Liu, D., et al. (2018). Serotonin receptor 2c-expressing cells in the ventral CA1 control attention via innervation of the Edinger–Westphal nucleus. Nat. Neurosci. 21, 1239–1250.

Loewy, A.D., and Saper, C.B. (1978). Edinger-Westphal nucleus: Projections to the brain stem and spinal cord in the cat. Brain Res. 150, 1–27.

Lovett-Barron, M., Andalman, A.S., Allen, W.E., Vesuna, S., Kauvar, I., Burns, V.M., and Deisseroth, K. (2017). Ancestral Circuits for the Coordinated Modulation of Brain State. Cell 171, 1411–1423.e17.

Maciewicz, R., Phipps, B.S., Grenier, J., and Poletti, C.E. (1984). Edinger-Westphal nucleus: cholecystokinin immunocytochemistry and projections to spinal cord and trigeminal nucleus in the cat. Brain Res. 299, 139–145.

McCullough, K.M., Morrison, F.G., Hartmann, J., Carlezon Jr, W.A., and Ressler, K.J. (2018). Quantified Coexpression Analysis of Central Amygdala Subpopulations. eNeuro 5, ENEURO.0010-18.2018.

Merighi, A. (2018). Costorage of High Molecular Weight Neurotransmitters in Large Dense Core Vesicles of Mammalian Neurons. Front. Cell. Neurosci. 12, 272.

Merighi, A., Salio, C., Ferrini, F., and Lossi, L. (2011). Neuromodulatory function of neuropeptides in the normal CNS. J. Chem. Neuroanat. 42, 276–287.

Moro, A., van Nifterick, A., Toonen, R.F., and Verhage, M. (2021). Dynamin controls neuropeptide secretion by organizing dense-core vesicle fusion sites. Sci. Adv. 7, eabf0659.

Nusbaum, M.P., and Blitz, D.M. (2012). Neuropeptide modulation of microcircuits. Curr. Opin. Neurobiol. 22, 592–601.

Nusbaum, M.P., Blitz, D.M., and Marder, E. (2017). Functional consequences of neuropeptide and small-molecule co-transmission. Nat. Rev. Neurosci. 18, 389.

Ogawa, M., Miyata, T., Nakajimat, K., Yagyu, K., Seike, M., Ikenaka, K., Yamamoto, H., and Mikoshibat, K. (1995). The reeler gene-associated antigen on cajal-retzius neurons is a crucial molecule for laminar organization of cortical neurons. Neuron 14, 899–912.

Okere, B., Xu, L., Roubos, E.W., Sonetti, D., and Kozicz, T. (2010). Restraint stress alters the secretory activity of neurons co-expressing urocortin-1, cocaine- and amphetamine-regulated transcript peptide and nesfatin-1 in the mouse Edinger–Westphal nucleus. Brain Res. 1317, 92–99.

Paxinos, G., and Franklin, K.B.J. (2019). Paxinos and Franklin’s the mouse brain in stereotaxic coordinates (Academic press).

Peirs, C., Williams, S.-P.G.P.G., Zhao, X., Walsh, C.E.E., Gedeon, J.Y.Y., Cagle, N.E.E., Goldring, A.C.C., Hioki, H., Liu, Z., Marell, P.S.S., et al. (2015). Dorsal Horn Circuits for Persistent Mechanical Pain. Neuron 87, 797–812.

Pérez-Escudero, A., Vicente-Page, J., Hinz, R.C., Arganda, S., and de Polavieja, G.G. (2014). idTracker: tracking individuals in a group by automatic identification of unmarked animals. Nat. Methods 11, 743–748.

Persoon, C.M., Hoogstraaten, R.I., Nassal, J.P., van Weering, J.R.T., Kaeser, P.S., Toonen, R.F., and Verhage, M. (2019). The RAB3-RIM Pathway Is Essential for the Release of Neuromodulators. Neuron 104, 1065–1080.e12.

Phipps, B.S., Maciewicz, R., Sandrew, B.B., Poletti, C.E., and Foote, W.E. (1983). Edinger-Westphal neurons that project to spinal cord contain substance P. Neurosci. Lett. 36, 125–131.

Piñol, R.A., Jameson, H., Popratiloff, A., Lee, N.H., and Mendelowitz, D. (2014). Visualization of oxytocin release that mediates paired pulse facilitation in hypothalamic pathways to brainstem autonomic neurons. PLoS One 9, e112138–e112138.

Van Den Pol, A.N. (2012). Neuropeptide transmission in brain circuits. Neuron 76, 98–115.

Pologruto, T.A., Sabatini, B.L., and Svoboda, K. (2003). ScanImage: flexible software for operating laser scanning microscopes. Biomed. Eng. Online 2, 13.

Pomrenze, M.B., Giovanetti, S.M., Maiya, R., Gordon, A.G., Kreeger, L.J., and Messing, R.O. (2019). Dissecting the Roles of GABA and Neuropeptides from Rat Central Amygdala CRF Neurons in Anxiety and Fear Learning. Cell Rep. 29, 13–21.e4.

Puntman, D.C., Arora, S., Farina, M., Toonen, R.F., and Verhage, M. (2021). Munc18-1 is essential for neuropeptide secretion in neurons. J. Neurosci. 41, 5980–5993.

Reardon, T.R., Murray, A.J., Turi, G.F., Wirblich, C., Croce, K.R., Schnell, M.J., Jessell, T.M., and Losonczy, A. (2016). Rabies Virus CVS-N2cΔG Strain Enhances Retrograde Synaptic Transfer and Neuronal Viability. Neuron 89, 711–724.

Rigney, N., Whylings, J., de Vries, G.J., and Petrulis, A. (2021). Sex Differences in the Control of Social Investigation and Anxiety by Vasopressin Cells of the Paraventricular Nucleus of the Hypothalamus. Neuroendocrinology 111, 521–535.

Romanov, R.A., Zeisel, A., Bakker, J., Girach, F., Hellysaz, A., Tomer, R., Alpár, A., Mulder, J., Clotman, F., Keimpema, E., et al. (2017). Molecular interrogation of hypothalamic organization reveals distinct dopamine neuronal subtypes. Nat. Neurosci. 20, 176–188.

Ryabinin, A.E., Cocking, D.L., and Kaur, S. (2013). Inhibition of VTA neurons activates the centrally projecting Edinger–Westphal nucleus: Evidence of a stress–reward link? J. Chem. Neuroanat. 54, 57–61.

Ryan, P.J., Ross, S.I., Campos, C.A., Derkach, V.A., and Palmiter, R.D. (2017). Oxytocin-receptor-expressing neurons in the parabrachial nucleus regulate fluid intake. Nat. Neurosci. 20, 1722– 1733.

Samuels, B.A., and Hen, R. (2011). Novelty-Suppressed Feeding in the Mouse. In Mood and Anxiety Related Phenotypes in Mice: Characterization Using Behavioral Tests, Volume II, T.D. Gould, ed. (Totowa, NJ: Humana Press), pp. 107–121.

Dos Santos Júnior, E.D., Da Silva, A. V., Da Silva, K.R.T., Haemmerle, C.A.S., Batagello, D.S., Da Silva, J.M., Lima, L.B., Da Silva, R.J., Diniz, G.B., Sita, L. V., et al. (2015). The centrally projecting Edinger–Westphal nucleus—I: Efferents in the rat brain. J. Chem. Neuroanat. 68, 22–38.

Schindelin, J., Arganda-Carreras, I., Frise, E., Kaynig, V., Longair, M., Pietzsch, T., Preibisch, S., Rueden, C., Saalfeld, S., Schmid, B., et al. (2012). Fiji: an open-source platform for biological-image analysis. Nat. Methods 9, 676–682.

Schöne, C., and Burdakov, D. (2012). Glutamate and GABA as rapid effectors of hypothalamic “peptidergic” neurons. Front. Behav. Neurosci. 6, 81.

Senn, V., Wolff, S.B.E., Herry, C., Grenier, F., Ehrlich, I., Gründemann, J., Fadok, J.P., Müller, C., Letzkus, J.J., and Lüthi, A. (2014). Long-Range Connectivity Defines Behavioral Specificity of Amygdala Neurons. Neuron 81, 428–437.

Siemian, J.N., Arenivar, M.A., Sarsfield, S., Borja, C.B., Erbaugh, L.J., Eagle, A.L., Robison, A.J., Leinninger, G., and Aponte, Y. (2021). An excitatory lateral hypothalamic circuit orchestrating pain behaviors in mice. Elife 10, e66446.

Silva, C., and McNaughton, N. (2019). Are periaqueductal gray and dorsal raphe the foundation of appetitive and aversive control? A comprehensive review. Prog. Neurobiol. 177, 33–72.

Silva, B.A., Mattucci, C., Krzywkowski, P., Murana, E., Illarionova, A., Grinevich, V., Canteras, N.S., Ragozzino, D., and Gross, C.T. (2013). Independent hypothalamic circuits for social and predator fear. Nat. Neurosci. 16, 1731–1733.

Skirboll, L., Hökfelt, T., Rehfeld, J., Cuello, A.C., and Dockray, G. (1982). Coexistence of substance P- and cholecystokinin-like immunoreactivity in neurons of the mesencephalic periaqueductal central gray. Neurosci. Lett. 28, 35–39.

Stanek, L.M. (2006). Cocaine- and amphetamine related transcript (CART) and anxiety. Peptides 27, 2005–2011.

Stuber, G.D., Hnasko, T.S., Britt, J.P., Edwards, R.H., and Bonci, A. (2010). Dopaminergic Terminals in the Nucleus Accumbens But Not the Dorsal Striatum Corelease Glutamate. J. Neurosci. 30, 8229 LP – 8233.

Sztainberg, Y., and Chen, A. (2012). Chapter 15 - Neuropeptide Regulation of Stress-Induced Behavior: Insights from the CRF/Urocortin Family. G. Fink, D.W. Pfaff, and J.E.B.T.-H. of N. Levine, eds. (San Diego: Academic Press), pp. 355–375.

Taniguchi, H., He, M., Wu, P., Kim, S., Paik, R., Sugino, K., Kvitsani, D., Fu, Y., Lu, J., Lin, Y., et al. (2011). A Resource of Cre Driver Lines for Genetic Targeting of GABAergic Neurons in Cerebral Cortex. Neuron 71, 995–1013.

Toyoshima, K., Kawana, E., and Sakai, H. (1980). On the neuronal origin of the afferents to the ciliary ganglion in cat. Brain Res. 185, 67–76.

Tritsch, N.X., Ding, J.B., and Sabatini, B.L. (2012). Dopaminergic neurons inhibit striatal output through non-canonical release of GABA. Nature 490, 262–266.

Ugolini, G. (2011). Chapter 10 - Rabies Virus as a Transneuronal Tracer of Neuronal Connections. In Research Advances in Rabies, A.C.B.T.-A. in V.R. Jackson, ed. (Academic Press), pp. 165– 202.

Vaaga, C.E., Borisovska, M., and Westbrook, G.L. (2014). Dual-transmitter neurons: functional implications of co-release and co-transmission. Curr. Opin. Neurobiol. 29, 25–32.

Vasconcelos, L.A.P., Donaldson, C., Sita, L. V., Casatti, C.A., Lotfi, C.F.P., Wang, L., Cadinouche, M.Z.A., Frigo, L., Elias, C.F., Lovejoy, D.A., et al. (2003). Urocortin in the central nervous system of a primate (Cebus apella): Sequencing, immunohistochemical, and hybridization histochemical characterization. J. Comp. Neurol. 463, 157–175.

Walf, A.A., and Frye, C.A. (2007). The use of the elevated plus maze as an assay of anxiety-related behavior in rodents. Nat. Protoc. 2, 322–328.

Wang, H.-L., Zhang, S., Qi, J., Wang, H., Cachope, R., Mejias-Aponte, C.A., Gomez, J.A., Mateo-Semidey, G.E., Beaudoin, G.M.J., Paladini, C.A., et al. (2019). Dorsal Raphe Dual Serotonin-Glutamate Neurons Drive Reward by Establishing Excitatory Synapses on VTA Mesoaccumbens Dopamine Neurons. Cell Rep. 26, 1128–1142.e7.

Wang, Q., Ding, S.-L., Li, Y., Royall, J., Feng, D., Lesnar, P., Graddis, N., Naeemi, M., Facer, B., Ho, A., et al. (2020). The Allen Mouse Brain Common Coordinate Framework: A 3D Reference Atlas. Cell 181, 936–953.e20.

Watabe-Uchida, M., Zhu, L., Ogawa, S.K., Vamanrao, A., and Uchida, N. (2012). Whole-Brain Mapping of Direct Inputs to Midbrain Dopamine Neurons. Neuron 74, 858–873.

Weitemier, A.Z., Tsivkovskaia, N.O., and Ryabinin, A.E. (2005). Urocortin 1 distribution in mouse brain is strain-dependent. Neuroscience 132, 729–740.

Wilson, T.D., Valdivia, S., Khan, A., Ahn, H.-S., Adke, A.P., Martinez Gonzalez, S., Sugimura, Y.K., and Carrasquillo, Y. (2019). Dual and Opposing Functions of the Central Amygdala in the Modulation of Pain. Cell Rep. 29, 332–346.e5.

Wu, M., Minkowicz, S., Dumrongprechachan, V., Hamilton, P., and Kozorovitskiy, Y. (2021a). Ketamine Rapidly Enhances Glutamate-Evoked Dendritic Spinogenesis in Medial Prefrontal Cortex Through Dopaminergic Mechanisms. Biol. Psychiatry 89, 1096–1105.

Wu, M., Minkowicz, S., Dumrongprechachan, V., Hamilton, P., Xiao, L., and Kozorovitskiy, Y. (2021b). Attenuated dopamine signaling after aversive learning is restored by ketamine to rescue escape actions. Elife 10, e64041.

Wu, Z., Autry, A.E., Bergan, J.F., Watabe-Uchida, M., and Dulac, C.G. (2014). Galanin neurons in the medial preoptic area govern parental behaviour. Nature 509, 325–330.

Xiao, L., Priest, M.F., Nasenbeny, J., Lu, T., and Kozorovitskiy, Y. (2017). Biased Oxytocinergic Modulation of Midbrain Dopamine Systems. Neuron 95, 368–384.e5.

Xiao, L., Priest, M.F., and Kozorovitskiy, Y. (2018). Oxytocin functions as a spatiotemporal filter for excitatory synaptic inputs to VTA dopamine neurons. Elife 7, e33892.

Xu, L., Bloem, B., Gaszner, B., Roubos, E.W., and Kozicz, T. (2009). Sex-specific effects of fasting on urocortin 1, cocaine- and amphetamine-regulated transcript peptide and nesfatin-1 expression in the rat Edinger–Westphal nucleus. Neuroscience 162, 1141–1149.

Yilmaz, M., and Meister, M. (2013). Rapid Innate Defensive Responses of Mice to Looming Visual Stimuli. Curr. Biol. 23, 2011–2015.

Zeisel, A., Hochgerner, H., Lönnerberg, P., Johnsson, A., Memic, F., van der Zwan, J., Häring, M., Braun, E., Borm, L.E., La Manno, G., et al. (2018). Molecular Architecture of the Mouse Nervous System. Cell 174, 999–1014.e22.

Zelikowsky, M., Hui, M., Karigo, T., Choe, A., Yang, B., Blanco, M.R., Beadle, K., Gradinaru, V., Deverman, B.E., and Anderson, D.J. (2018). The neuropeptide Tac2 controls a distributed brain state induced by chronic social isolation stress. Cell 173, 1265–1279.

Zhang, L., Hernández, V.S., Zetter, M.A., and Eiden, L.E. (2020). VGLUT-VGAT expression delineates functionally specialised populations of vasopressin-containing neurones including a glutamatergic perforant path-projecting cell group to the hippocampus in rat and mouse brain. J. Neuroendocrinol. 32, e12831.

Zhang, S., Qi, J., Li, X., Wang, H.-L., Britt, J.P., Hoffman, A.F., Bonci, A., Lupica, C.R., and Morales, M. (2015). Dopaminergic and glutamatergic microdomains in a subset of rodent mesoaccumbens axons. Nat. Neurosci. 18, 386–392.

Zhang, Z., Zhong, P., Hu, F., Barger, Z., Ren, Y., Ding, X., Li, S., Weber, F., Chung, S., Palmiter, R.D., et al. (2019). An Excitatory Circuit in the Perioculomotor Midbrain for Non-REM Sleep Control. Cell 177, 1293–1307.e16.

Zhou, M., Liu, Z., Melin, M.D., Ng, Y.H., Xu, W., and Südhof, T.C. (2018). A central amygdala to zona incerta projection is required for acquisition and remote recall of conditioned fear memory. Nat. Neurosci. 21, 1515–1519.

